# Bifurcation drives the evolution of assembly-line biosynthesis

**DOI:** 10.1101/2021.06.23.449585

**Authors:** Thomas J. Booth, Kenan A. J. Bozhüyük, Jonathon D. Liston, Ernest Lacey, Barrie Wilkinson

## Abstract

Reprogramming biosynthetic assembly-lines is a topic of intense interest. This is unsurprising as the scaffolds of most antibiotics in current clinical use are produced by such pathways. The modular nature of assembly-lines provides a direct relationship between the sequence of enzymatic domains and the chemical structure of the product, but rational reprogramming efforts have been met with limited success. To gain greater insight into the design process, we wanted to examine how Nature creates assembly-lines and searched for biosynthetic pathways that might represent evolutionary transitions. By examining the biosynthesis of the anti-tubercular wollamides, we show how whole gene duplication and neofunctionalization can result in pathway bifurcation. Importantly, we show that neofunctionalization occurs primarily through intragenomic recombination. This pathway bifurcation leads to redundancy, providing the genetic robustness required to enable large structural changes during the evolution of antibiotic structures. Should the new product be none-functional, gene loss can restore the original genotype. However, if the new product confers an advantage, depreciation and eventual loss of the original gene creates a new linear pathway. This provides the blind watchmaker equivalent to the ‘design, build, test’ cycle of synthetic biology.

## Introduction

Microbial natural products produced by modular biosynthetic assembly-lines, i.e. (type I) polyketide synthases (PKSs)^1^ and non-ribosomal peptide synthetases (NRPSs)^2^, are an important source of pharmaceutical and agrochemical agents. Examples include well known antibacterial molecules such as the polyketide insecticide spinosyn^3^ and NRPS derived penicillins^4^. Importantly, biosynthetic assembly-lines provide thousands of natural product scaffolds, including many of our essential clinical agents.

Essentially, NRPS and PKS modular megasynth(et)ases give rise to highly functionalised biopolymers from a broad variety of monomers, referred to as extender units. Hundreds of extender units have been reported, typically derived from malonate in the case of PKSs^5^ or amino acids in the case of NRPSs^6,7^. They are likened to assembly-line processes due to their hierarchical and modular structures in which multiple, repeating modules of enzymatic domains catalyse the incorporation of an extender unit into the growing chain, along with any programmed additional chemical modifications, before transferring the elongated chain to the next module. The archetypical minimal assembly-line module consists of three ‘core’ domains. Firstly, a domain for the selection and activation of an extender unit, the acyltransferase (AT) domains for PKSs or adenylation (A) domains for NRPSs. The activated substrate is then covalently attached to a prosthetic phosphopantetheine group of a small acyl carrier protein (ACP; PKSs) or peptidyl carrier protein (PCP; NRPSs) domain. Finally, the ketosynthase (KS; PKSs) or condensation (C; NRPSs) domains then link the covalently bound substrates to the growing polyketide or peptide chain. Although the exact mechanisms and ancillary domains of PKSs and NRPSs differ, the fundamental principal is that modules condense covalently bound substrates in a linear fashion. The inherent logic of this mechanism means that there is a direct relationship between the sequence of domains in an assembly line and the chemical structure of the resulting molecule^1,8,9^. In principle this relationship enables the prediction of natural product chemical structures directly from DNA sequences. In turn, this logic has inspired efforts to rationally reprogramme assembly-lines to produce tailor-made molecules.

Numerous examples of assembly line engineering have been reported, however many display productivities well below that of the parental (wild type) system. Insights into structural flexibility, inter-domain communication, and the role of proof-reading by catalytic domains preempted novel strategies to engineer assembly-line proteins^10–14^. There is also an increasing body of evidence suggesting that we might further improve our ability to engineer these systems if we had a better understanding of their evolution^1,15–18^.

With this latter point in mind, we have been searching for biosynthetic pathways that may represent transitional evolutionary states and provide exemplar systems to inform future work. Natural selection acts upon phenotypes, yet even a small change to the structure of a natural product can have profound effects on bioactivity and, thus, the fitness of the producing organism. Therefore, we hypothesised that strains encoding BGCs evolving new functionalities might be expected to produce multiple related products (congeners). Here, we describe the BGC encoding the wollamide-desotamide family of antibiotics, which represents an evolutionary snapshot of a modular NRPS assembly line.

The wollamides are cyclic hexapeptides (Fig. 1a) first reported in 2014^19^. The only known producer of the wollamides is *Streptomyces* sp. MST-115088^19^ and they exhibit potent antimycobacterial activity that has attracted the attention of synthetic medicinal chemists^20–22^. Along with the wollamides, *Streptomyces* sp. MST-115088 produces a related group of hexapeptides, the desotamides^23^. The desotamides and wollamides share a common peptide scaffold, except for a single residue change from glycine in the desotamides to D-ornithine in the wollamides (Fig. 1a). It is important to note that the previously identified desotamide producer, *Streptomyces scopuliridis* SCSIO ZJ46, is not reported to produce wollamides and the desotamide (*dsa*) BGC follows canonical NRPS logic^24^. The NRPS is encoded by 3 genes (*dsaI, dsaH* and *dsaG*) encoding two modules each. Importantly, module six, encoded by DsaG, incorporates glycine as the final amino acid. The ability of *Streptomyces* sp. MST-115088 to produce congeners with D-ornithine and glycine in the same position is therefore difficult to rationalise as, under a canonical NRPS mechanism, A-domains are responsible for the selection and activation of specific amino acid substrates. While A-domains can activate structurally related amino acids giving rise to families of structurally similar congeners (for example the combinations of valine, leucine, or *allo*-leucine at positions 3 and 4 of the wollamides-desotamides), the ability of an A-domain to activate substrates with such varying physico-chemical properties as glycine and ornithine is without precedent. Therefore, to explain the production of wollamides and desotamides by a single strain we hypothesised three scenarios (Fig. 1b): dual specificity of module six for glycine and ornithine; duplicated genes encoding module six homologues, each specific to glycine or ornithine respectively; or duplicated BGCs where, as above, the final modules of each BGC are specific to glycine or ornithine.

**Fig. 1.**
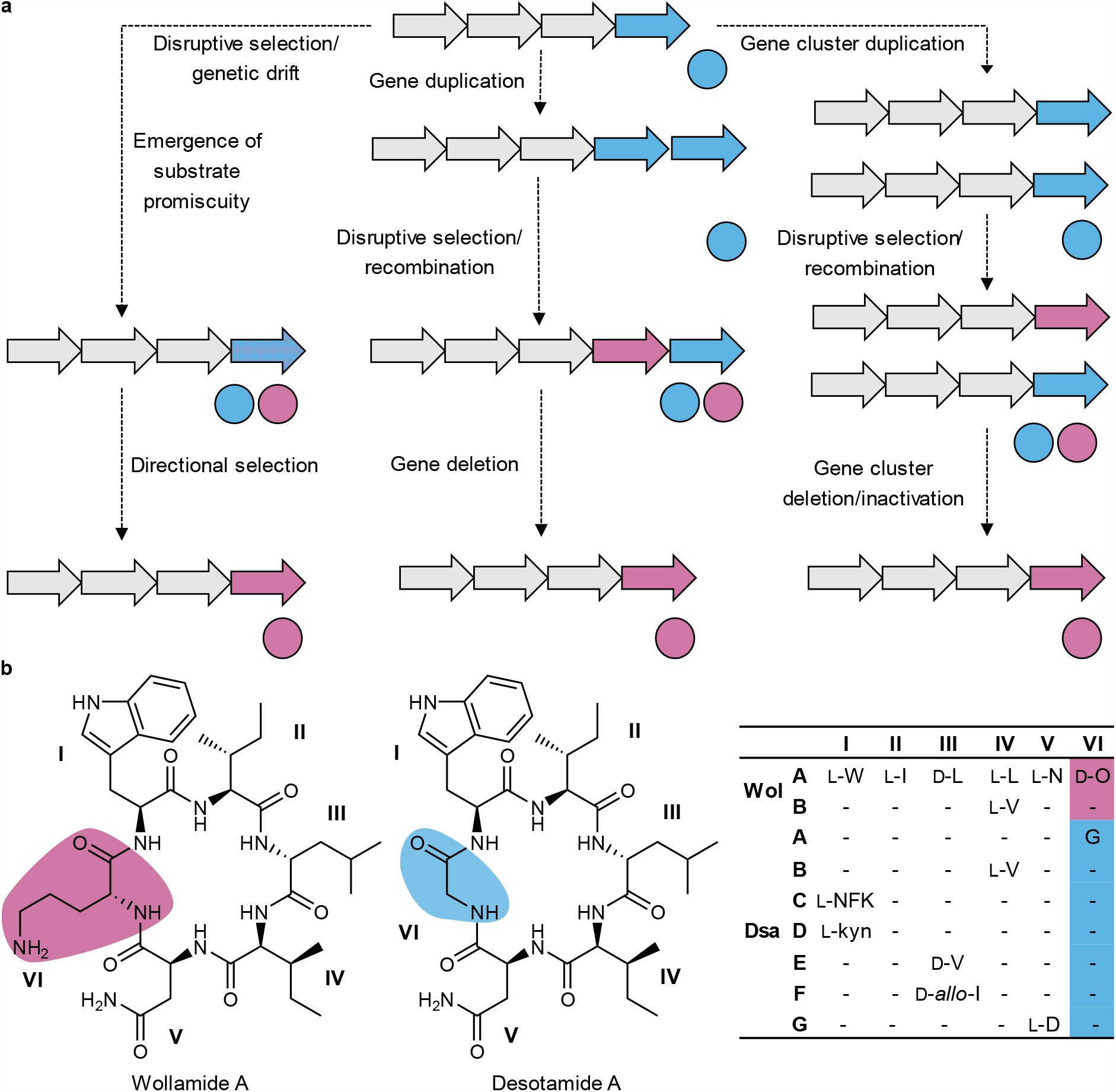
Intermediate chemotypes in the evolution of natural product BGCs. **a**, a cartoon demonstrating the three main models of gene cluster evolution depicting the transition of a fictitious gene cluster from chemotype A to chemotype B (coloured circles). Evolutionary processes are represented by dashed lines. **b**, The structural diversity of the wollamides and desotamides. The D-ornithine and glycine, which define the wollamides and desotamides respectively, are highlighted. The table on the right-hand side shows the variable positions of the various wollamide and desotamide congeners (NFK = *N*-formyl kyunerine).

Herein, we describe our combined bioinformatics, *in vivo* engineering, and biochemical analysis demonstrating how *dsa* evolved from an ancestral wollamide producing BGC via the process of gene duplication and intragenomic recombination with a second NRPS encoding locus. This allows us to propose an update to the current model for the evolution of assembly line BGCs through which duplication and the resulting bifurcation reduces selective pressure and drives the evolution of new functions.

## Results

### Desotamides and wollamides are produced by a bifurcated NRPS assembly-line

The genome of *Streptomyces* sp. MST-115088 was sequenced using Pacific Biosciences RS2^25^ single molecule technology and assembled using the HGAP3^26^ pipeline to generate a single 7.9 Mb chromosomal assembly (Supplementary Table 1)(GenBank:CP074380). Analysis of the assembly using antiSMASH v4.0^8^ allowed us to rapidly identify the wollamide (*wol*) BGC (Supplementary Table 2), which was then compared to the desotamide (*dsa*) BGC previously reported from *Streptomyces scopuliridis* SCSIO ZJ46 (GenBank: KP769807) (Fig. 2ab). Five additional desotamide producers (MST-70754, MST-71458, MST-71321, MST-94754 and MST-127221) were identified through metabolomic analysis of Microbial Screening Technologies unique collection of more than fifty thousand Australian actinomycete strains (Supplementary Information). The desotamide BGC (Supplementary Tables 3-7) of these strains display functionally identical architectures (GenBank: MZ093610-MZ093614) to that reported from *Streptomyces scopuliridis* SCSIO ZJ46^24^ (>95% nucleotide identity) (Supplementary Fig. 1). The *wol* BGC displays a similar architecture to the *dsa* BGCs but contains two genes *wolG1* and wolG2 that are duplicates of the *dsaG* gene which encodes the final two modules of the NRPS. Similarly, there are duplicated homologues of the *dsaF* gene (*wolF1* and *wolF2*), which encodes an MbtH-like protein involved in A-domain functionality^27,28^. The *wol* BGC also contains six additional genes *wolRSTUVW* predicted to be involved in the biosynthesis of L-ornithine consistent with the presence of ornithine in position 6 of the wollamides.

**Fig. 2.**
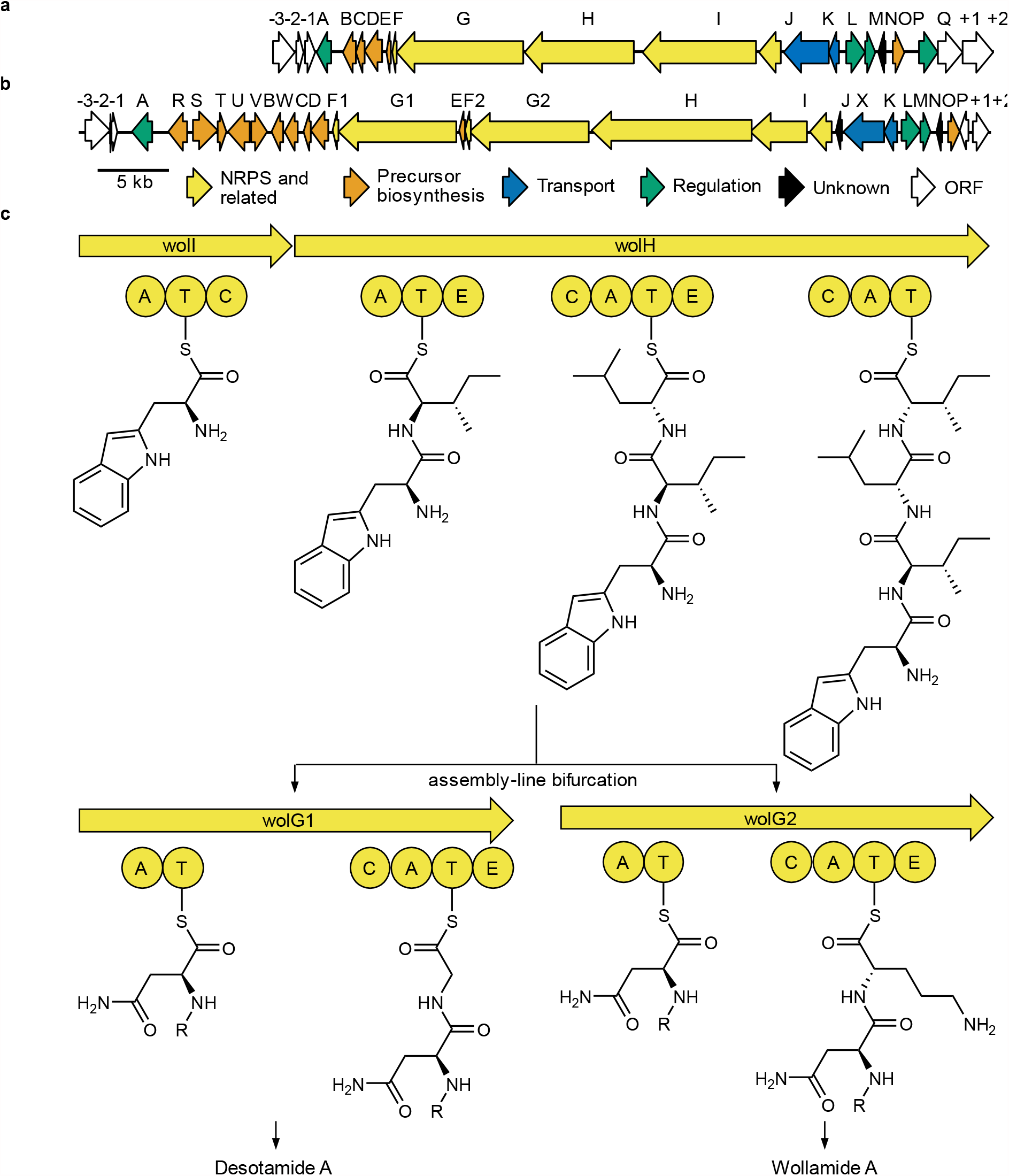
The genetic basis for bifurcated biosynthesis of desotamide and wollamide by *Streptomyces*. sp. MST-110588. **a**, The architecture of the desotamide producing *dsa* BGC from *Streptomyces scopuliridis* SCSIO ZJ4624. **b**, The architecture of the wollamide and desotamide producing *wol* BGC from *Streptomyces* sp. MST110588. **c**, The proposed biosynthetic pathway of the wollamides and desotamides. The first three condensations are catalysed by WolIH. The final two condensations are catalysed by WolG1 to produce desotamides or WolG2 to produce wollamides.

During desotamide biosynthesis, DsaG is responsible for the final two rounds of peptide chain elongation. We hypothesised, therefore, that WolG1 and WolG2 may encode two forks of a bifurcated biosynthetic pathway where the first four rounds of peptide elongation proceed via the colinear activity of WolI and WoH, but the final two elongation steps are catalysed either by WolG1 or WolG2, yielding desotamide or wollamide products respectively (Fig. 2c). Consistent with this hypothesis, *in silico* analysis of A-domain specificities^29,30^ predicted the substrates for the final A-domains of WolG1 and WolG2 (henceforth WolG1A2 and WolG2A2) to be glycine and L-ornithine (Supplementary Table 8).

### Engineering wollamide production in a desotamide-only producing strain

To confirm our biosynthetic hypothesis and to explore the evolutionary relationship between the wollamide and desotamide pathways, we sought to engineer the co-production of wollamides into a desotamide-only producing strain through the heterologous expression of *wolG2*.

As NRPS genes are typically large (for reference, *wolG2* is 7.9 kb) and difficult to clone through conventional methods we generated pBO1 (Supplementary Fig. 2), a *Streptomyces* expression vector capable of propagating in both *E. coli* and *S. cerevisiae*. This allows larger genes to be assembled efficiently in yeast via transformation associated homologous recombination^31^ downstream of a constitutive promotor and subsequently transferred from *E. coli* to *Streptomyces* spp. through conjugal transfer (Supplementary Information). To assemble a *wolG2* expression plasmid, pBO1 along with target fragments amplified from genomic DNA were transformed into *S. cerevisae* CEN.PK 2-1C^32^. The resulting plasmid pBO1_*wolG2* was transformed into the desotamide producer *Streptomyces* sp. MST-70754^33^ via conjugation.

*Streptomyces* sp. MST-70754 and progeny carrying pBO1_*wolG2* were grown under desotamide producing conditions and methanolic culture extracts were then analysed by LCMS in comparison to the wollamide producer *Streptomyces* sp. MST-115088. The presence of both wollamide and desotamide congeners was confirmed for the engineered strains by comparison of retention time, isotopic masses and MS/MS fragmentation of the compounds produced by the native wollamide producer (desotamide: [M+H]^+^ = 697.4047, [M+Na]^+^ = 719.382; wollamide A [M+H]^+^ 754.4655, [M+Na]^+^ = 776.4419) (Fig. 3) (Supplementary Fig. 3).

**Fig. 3.**
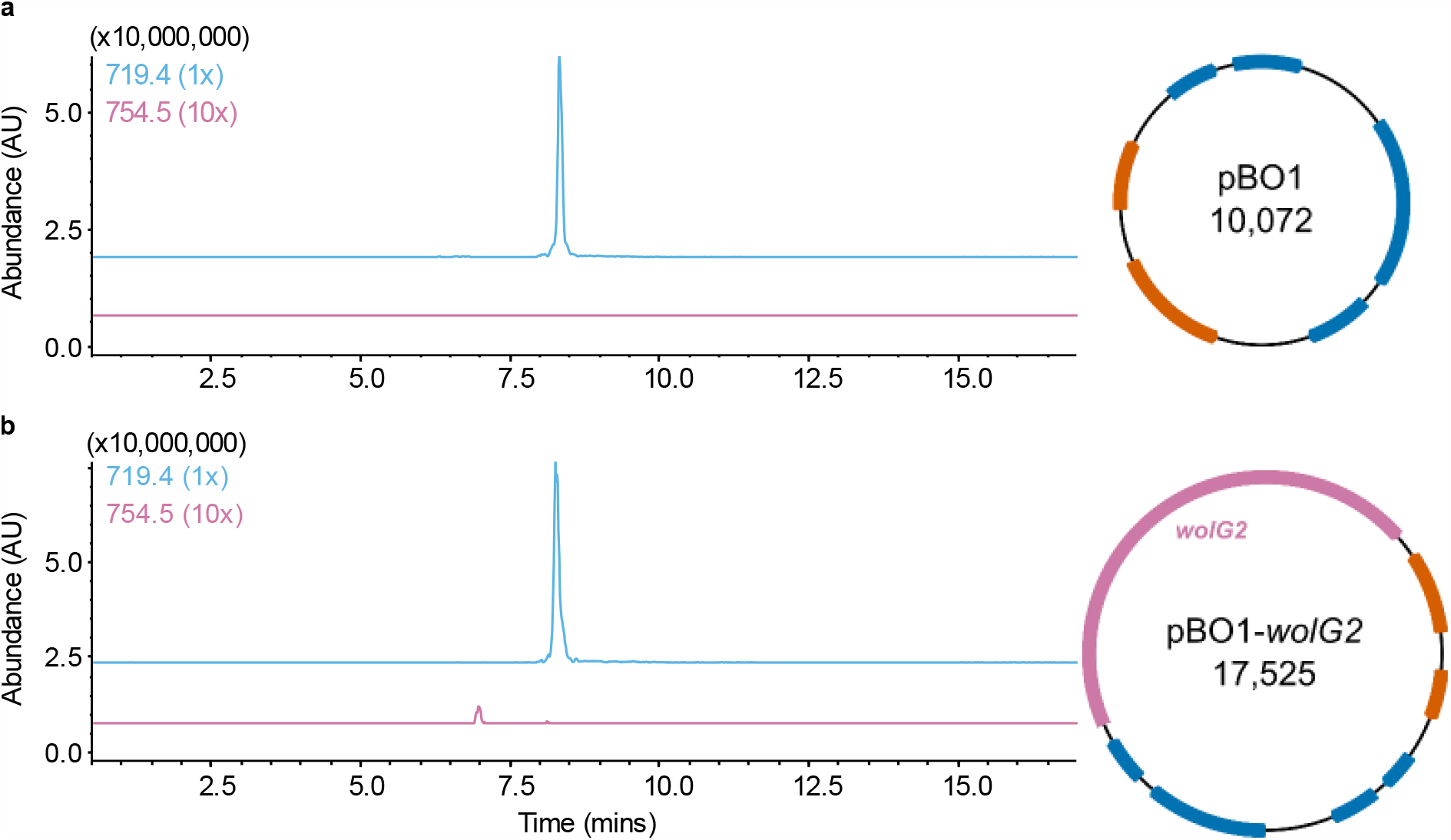
Heterologous production of the wollamides by overexpression of *wolG2*. **a**, The extracted ion chromatograms (EIC) for masses corresponding to wollamide A ([M+H] = 754.4) and desotamide A ([M+H] = 719.4) from wild type *Streptomyces* sp. MST-70754 transformed with the plasmid pBO1, producing only desotamide A. **b**, The EICs of wollamide A and desotamide A from *S*. sp. MST-70754 transformed with the plasmid pBO1-wolG2, showing production of both the wollamides and desotamides.

This engineering strategy relied on the assumption that the docking domains (DDs) of DsaH and WolG2 that mediate NRPS interactions would still function as a pair^34,35^. While the production of wollamides shows that the interaction is still possible, the low relative yields of wollamides versus desotamides suggested that some depreciation has occurred.

To investigate this, we compared the *N*-terminal DD primary sequences of WolG1 and WolG2 (Supplementary Fig. 4), as well as calculated homology models of the DDs in complex with the WolH DD (Supplementary Fig. 5). Alignment of the DDs revealed a glutamate to alanine mutation at position (E16A). Homology modelling with the previously characterised PaxB docking domain (6TRP_1)^36^ revealed that the E16A amino acid change in WolG2 leads to the loss of a salt bridge formed with R3504 from WolH, most likely causing decreased DD pair affinities, consistent with the observed differences in titers.

### Evolution of adenylation domain specificity via intragenomic recombination

The unusual architecture of the *wol* BGC led us to consider potential mechanisms for its evolution. The high similarity of *wolG1* to *wolG2* (80.9%) and *wolG1*/*wolG2* to *dsaG* (74.3%71.2% respectively) is indicative of an ancestral gene duplication event^37^. However, despite this high similarity, there is a notable drop in nucleotide identity within the region coding for the final adenylation domains which is also manifest in the gene products (54.6% nucleotide and 29.2% protein sequence similarity) (Supplementary Fig. 6). Given the overall similarity of these genes we deemed it unlikely that such high sequence variation could emerge through speciation (point mutation) alone. Furthermore, phylogenetic reconstructions of A-domains hinted at independent evolutionary histories when compared to the rest of the assembly line (Supplementary Figures 7-9). Similar patterns have been observed in other bacterial NRPS clusters^18,38–40^

Many studies have highlighted the role recombination plays in the evolution of assembly-lines^1,41^. More specifically, given the reduced rate of horizontal gene transfer between distantly related taxa and the high rate of heterogeneity between recombinant sequences has, it has been speculated that intragenomic recombination within ancestral strains can provide opportunities for assembly-line diversification ^18,42^. To assess the possible role of intragenomic recombination in the evolution of the *wol* BGC we generated a nucleotide sequence alignment of all thirty-four NRPS-associated adenylation domains present in the *Streptomyces* sp. MST110588 genome for analysis using the Recombination Detection Programme 4 (RDP4)^43^.

Using our dataset, RDP4^43^ predicted 29 potential intragenomic recombination events (Supplementary Fig. 10 and Supplementary Table 9), 11 of which were supported by two or more methods, allowing recombination breakpoints to be predicted. Only two recombination events were supported unanimously, including one event, in which *wolG1A2* was identified as a recombinant sequence with *wolG2A2* as the major parent and an adenylation domain encoding sequence from elsewhere in the genome, encoded by *orf6595A*, as the minor parent (Fig 4a-c and Supplementary Table 9). The predicted recombinant region lies between the conserved motif A2 (*N*-terminal) and motifs A5 and A6 (*C*-terminal) of the A-domain, thus comprising the flavodoxin subdomain^18,38,40^ and a large portion of the *N*-terminal subdomain (Fig. 4d). Crucially, this would allow for the substitution of the amino acid binding pocket and catalytic P-loop while maintaining the C-A -domain interface (Supplementary Fig. 11). This pattern is seen frequently in our dataset (Supplementary Fig. 10) and may suggest an advantage for maintaining the structural relationships between the P-loop and substrate binding pocket.

**Fig. 4.**
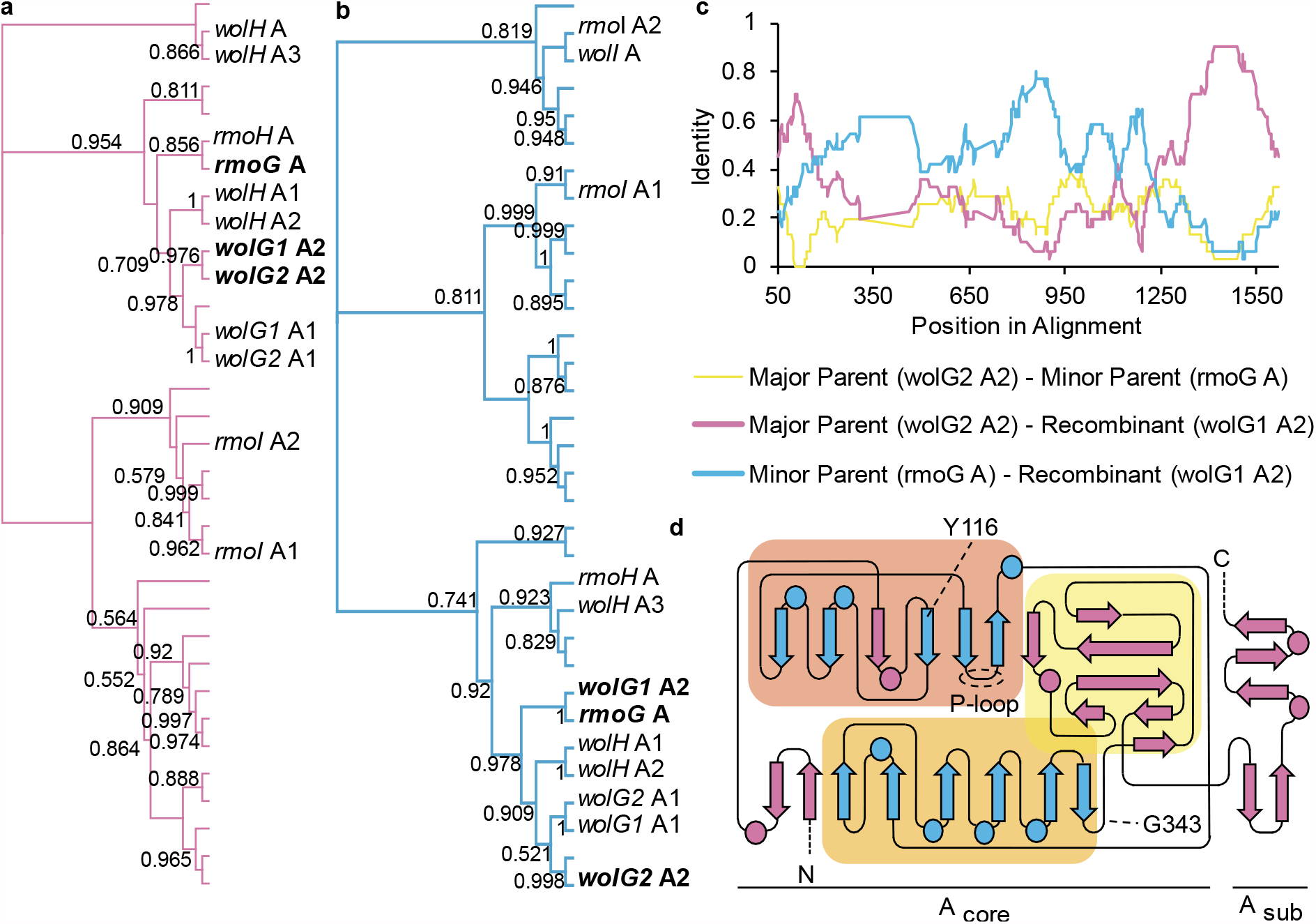
Evidence for the intragenomic recombination of adenylation domains. **a**, Phylogeny of the external region of the adenylation domains, showing the close relationship of the major parent, *wolG2* A2, and the recombinant, *wolG1* A2. **b**, Phylogeny of the internal region of the adenylation domains, showing the close relationship of the minor parent, *rmoG* A, and the recombinant, *wolG1* A2. **c**, Identity between the three adenylation domains. **d**, Topology of an adenylation domain adapted from Conti et al.74, showing the predicted recombination sites Y116 and G343. Features are colour-coded according to Fig 4a.

Importantly, the adenylation domain encoded by *orf6595* was predicted *in silico* to select for glycine (Supplementary Table 8) meaning that the predicted recombination event could theoretically convert a wollamide producing assembly line into a desotamide producer.

### Biochemical validation of A-domain substrate specificities

To gain insight into the function of the minor parent BGC, the sequences of Orf6595 and proteins encoded by the surrounding genes were searched against the MIBiG database^44^ using cblaster^45^. The search identified the BGC as homologous to the rimosamide (*rmo*) BGC from *Streptomyces rimosus* ATCC 10970^46^ (Supplementary Table 10). More specifically, Orf6595 is a homologue of RmoG which encodes a single NRPS module (C-A-T) that is known to incorporate glycine into the rimosamide peptide chain. This was consistent with bioinformatic predictions of the Orf6595 adenylation domain active site providing strong evidence that recombination between the adenylation domains of *orf6595* and *wolG2* could confer specificity to glycine (Supplementary Fig. 12).

To verify this prediction and examine the substrate specificity of all of the A-domains of interest, pET28a hexa-histidine tagged WolG1A2, WolG2A2 and Orf6595A constructs were cloned for expression based upon the A-domain boundaries as described in Crüsemann et al^38^. These were expressed in *E. coli* Rosetta 2(DE3)pLysS and purified using Ni-Affinity chromatography. Initially, the resulting protein was insoluble, however co-expression with the MbtH-like protein WolF2 (expressed from pCDFDuet-1) enabled the purification of soluble protein in each case. The ability of these isolated A-domains to activate each of the twenty proteinogenic amino acids and L-ornithine was then measured using an hydroxylamine trapping assay^47^ (Supplementary Table 11). WolG2A2 adenylates L-ornithine, in line with our hypothesis (Fig. 5a); however, it was also capable of activating other substrates albeit with lower efficiency. Most noticeably, WolG2A2 accepted L-aspartate (58% activity relative to L-ornithine) and L-asparagine (44% activity relative to L-ornithine) as substrates, but wollamide analogues in which aspartate or asparagine were substituted for ornithine were not identified in culture extracts of *Streptomyces* sp. MST-110588 despite targeted LCMS analysis. In contrast, both WolG1A2 and Orf6595A activate glycine in a highly specific manner (Fig. 5a). These data corroborate our hypothesis that an historic recombination with *orf6595* could alter the substrate specificity of the module six adenylation domain of WolG2 from L-ornithine to glycine. To test this hypothesis further, we produced a hybrid A-domain encoding gene sequence representing the hypothetical ancestral recombinant, based on *wolG2A2* and *orf6595* sequences (Fig. 5b). The resulting gene product was purified and assayed as above and found to be highly selective for glycine (Fig. 5a).

**Fig. 5.**
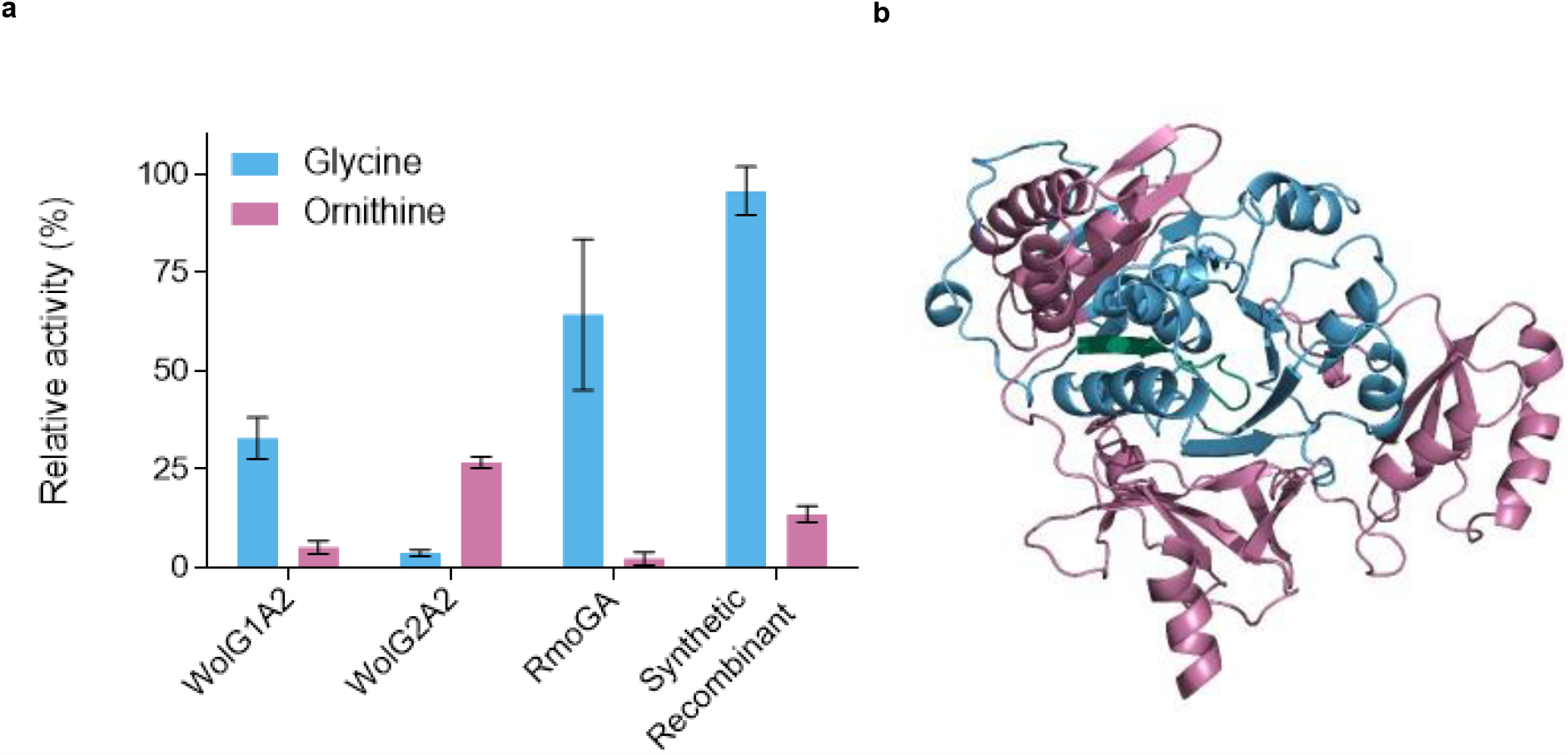
Substrate specificities of wild type and recombinant adenylation domains. **a**, Relative activities of adenylation domains as determined by hydroxylamine trapping assays47. **b**, Structural homology model of the synthetic recombinant adenylation domain. The model is colour coded depending on the origin of the peptide sequence, pink for WolG2A2 and blue for RmoGA. The P-loop is highlighted in green.

Taken together with combined genomic, *in silico* and *in vivo* data above, these biochemical data show how the contemporary desotamide BGCs have evolved from an ancestral wollamide-like BGC through the process of gene duplication, intragenomic recombination and gene loss.

## Discussion

How NRPS and PKS assembly line biosynthetic pathways evolve is a question of perennial interest in natural products research. An understanding of this process offers new avenues for developing rational approaches for bioengineering and the targeted production of new molecules. Evolutionary analyses of publicly available BGC sequences are common in the literature^18,48,49^, but it is unprecedented to find an example of a BGC that represents a snapshot of assembly line evolution. In this work, we studied *Streptomyces* sp. MST110588, a co-producer of desotamide and wollamide hexapeptide antibiotics. Despite their close structural relationship, it was not obvious how such a mixture of congeners might be assembled based on canonical NRPS function, pointing to an unusual biosynthetic pathway.

The architecture of the wollamide BGC is highly similar to the previously described desotamide BGC, however a striking difference is that it contains duplicates (*wolG1* and *wolG2*) of *dsaG* which encodes the final NRPS protein of the desotamide assembly line. This suggested a bifurcated assembly-line was responsible for the observed mixture of desotamide and wollamide congeners (Fig. 2c). In this scenario, WolIH is responsible for condensation of the first four amino acids prior to extension with either WolG1, producing desotamides, or WolG2, producing wollamides. This hypothesis was supported by bioinformatic (Fig. 4) and biochemical analysis (Fig. 5a) of the A-domains WolG1A2 and WolG2A2, which encode for selection and activation of glycine and L-ornithine respectively. Further *in vivo* evidence came from strain engineering in which expression of WolG2 in a desotamide-only producing strain led to biosynthesis of additional wollamide congeners (Fig. 3).

Comparison of the protein sequences of the NRPSs encoded by *wolG1*/*G2* indicated a high degree of sequence conservation except for the region coding for the final adenylation domains. These data were indicative of a recombination event rather than of divergence through speciation alone, and we subsequently analysed all 34 A-domain sequences present in the *Streptomyces* sp. MST110588 genome using the recombination detection program RDP4^43^. This identified *wolG2* as the major parent, and an NRPS gene *orf6595* (encoded approximately 3 Mb away on the chromosome) as the minor parent, of a recombination event that formed *wolG1*. Using the cblaster tool we identified the BGC containing *orf6595* as a homologue of the rimosamide BGC^46^. Orf6595 is a homologue of RmoG that selects for glycine. This was verified by subsequent bioinformatics and biochemical analysis. To validate the recombination event predicted by RDP4 we used the contemporary sequences of *wolG2A2* and *orf6595* to recapitulate the predicted ancestor and generated a synthetic A-domain. Subsequent biochemical analysis of the isolated protein showed it was selective for glycine activation as predicted.

Based on these combined data we can confidently trace the evolutionary history of the wollamide and desotamide BGCs (Fig. 6a). First, a gene duplication event in an ancestral wollamide (or wollamide-like) BGC resulted in a redundant copy of the bimodular NRPS encoding modules 5 and 6 of the assembly line. Subsequently, an intragenomic recombination event between the DNA encoding the adenylation domains of the duplicated module six in the *wolG* homologue and *orf6595* resulted in an intermediate NRPS selective for glycine. This progenitor is the ancestor of all wollamide and desotamide producing strains which, through additional selection and mutations, became the lineage of the *wol* BGC observed in the contemporary genome of *Streptomyces* sp. MST110588 capable of bifurcated biosynthesis. In a divergent lineage, the ancestral gene encoding the L-ornithine-specific adenylation domain along with the duplicated MbtH-like protein encoding gene and associated genes encoding L-ornithine biosynthesis were lost through gene deletion, ultimately resulting in the contemporary *dsa* BGCs producing only desotamides. These observations are consistent with the presence of a redundant epimerase domain located in the second modules of WolG1 and all DsaG homologues^50,51^. Moreover, in *Streptomyces* MST110588 production of wollamide congeners was more than an order of magnitude lower than that of desotamides. Our engineered strain of the desotamide producer, *Streptomyces* sp. MST-70754, expressing *wolG2* also showed this large difference in titer. This observation suggested that the deprecation of interactions between the C-and N-terminal docking domains of WolH/DsaH and WolG2 respectively. To assess this, we calculated homology models which revealed that a key E16A amino acid change in WolG2 leads to the loss of a salt bridge formed with R3504 from WolH, most likely causing decreased DD pair affinities, explaining the observed differences in peptide titers. The depreciation of this interaction is consistent with a loss of selective pressure for wollamide production and may indicate drift towards a desotamide-only BGC.

**Fig. 6.**
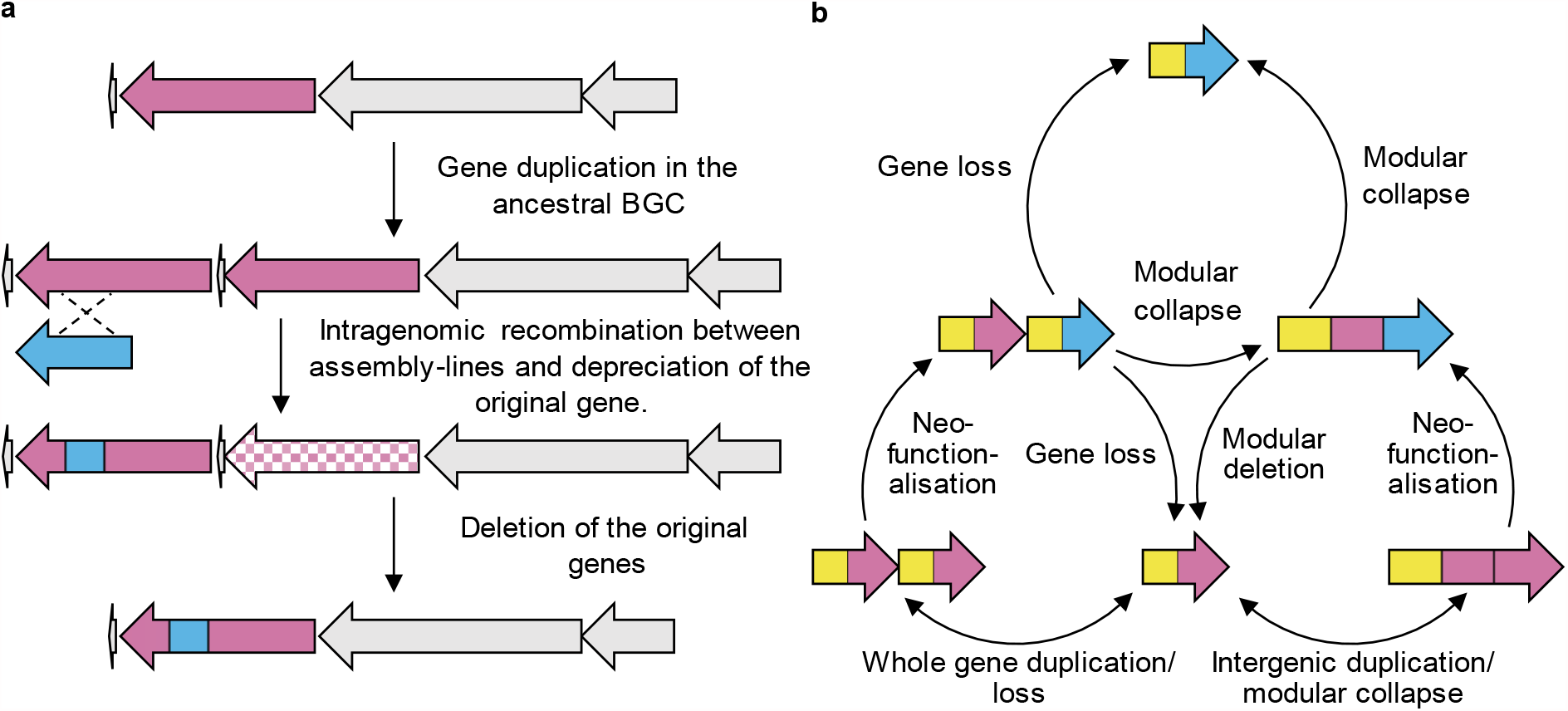
The evolution of biosynthetic assembly-lines. **a**, Proposed evolutionary history of the wollamide (*wol*) and desotamide (*dsa*) BGCs. **b**, An updated model for the evolution of biosynthetic assembly-lines.

Gene duplication and divergence is a central mechanism behind the evolution of new gene functions^37,52,53^. Gene duplications may occur as whole gene duplications (characterised by the duplication of an entire gene), or as partial or intragenic duplications (characterised by incomplete duplication of the ancestral gene and can result in attenuated or lengthened duplicates respectively^37^). There is convincing evidence for the role of intragenic duplications during assembly line evolution (Fig. 6b). Such processes are often cited as the origin of multi-modularity^1,54^ and there is significant evidence for this process, especially in PKS assembly lines. In contrast, whole gene duplication has the potential to create bifurcated biosynthetic pathways, as evidenced here by the *wol* BGC, or parallel pathways, as evidenced by the recently published BE-18257A-C and pentaminomycins A-E BGC^55^. Through either mechanism, whole gene duplication creates redundancy and has the potential to reduce selective pressure for maintenance on both alleles^37,53^. In this fashion, a gene may evolve new functionality without losing the original product. Thus, neofunctionalization of one copy becomes possible without strong selective pressure against the loss of a phenotype and may follow a more gradual route for the emergence of a new activity. Subsequently, the original allele can be lost if there is no advantage and selection pressure for its maintenance leading to a new linear pathway (Fig. 6b). Moreover, this model provides a scenario where rounds of duplication and neofunctionalization can occur but if no advantage is gained then the duplicated allele can be lost, restoring the original genotype. As genes and pathways are subject to multiple selective forces and it is unclear how common this mechanism may be in Nature. This is compounded by the homogenising effects of gene conversion/concerted evolution that can make it difficult to reliably distinguish duplication events^1,54,56,57^. However, the *wol* BGC provides compelling evidence that this process has occurred at least once, during the evolution of desotamide biosynthesis.

Finally, the question arises as to whether our knowledge of the NRPS evolution can also be used to develop new engineering approaches. Recently, several efficient and/or highly productive engineering strategies have been published^58–61^. Yet efficient engineering of these often-huge biosynthetic machineries to produce novel bioactive NRPs is an ongoing challenge. Understanding the evolution of these multifunctional enzymes might provide new insight for engineering and discovering new peptide based therapeutic agents^62^. A key feature of desotamide evolution that could be exploited for future NRPS engineering efforts is the intragenomic recombination between distinct NRPS encoding BGCs. The recombination event we predicted takes place within the boundaries of A domains, consistent with previous^38,39,63^ and more recent work^18,40^. More specifically, the predicted recombination allows for the substitution of the substrate specificity conferring active site and the ATP/phosphate binding catalytic P-loop^64^ while functionally maintaining most of the A-T^65^ and all of the C-A interdomain contacts^66–68^. This pattern is frequently seen in our dataset, suggesting an evolutionary advantage for maintaining the structural relationships between the P-loop and substrate binding pocket. Our results are supported by two other independent studies, further highlighting its significance^18,40^.

Calcott and co-workers analysis of recombination hotspots within C-A-T tri-domains (modules) from *Pseudomonas, Bacillus* and *Streptomyces* species identified the same recombination sites as predicted for the formation of *wolG1A2*^40^. Subsequently, Baunach and co-workers systematic *in silico* dissection of many individual recombination events unveiled the striking commonality of A domain recombination events in nature^18^. Specifically, these recombination events target variable portions of the Acore domains to modulate A domain substrates while domain-domain interactions and the flexible Asub domain largely remained unaffected. These studies, taken together with the evolutionary and biochemical evidence presented here, must inevitably lead to a change in paradigm of future NRPS engineering experiments. Established engineering principles are being overhauled by the increasing evidence base. In particular, the idea that C and A domains have coevolved^69,70^, resulting in strong acceptor site specificity of C domains^71,72^ is in question^18,40,60^. This hypothesis is not congruent with the observation that A domain recombination are responsible for the diversification of many NRPs^18,38,73^, not least the L-ornithine to glycine change demonstrated here. Such diversification would be impossible if C domain ‘gatekeeping’ was universal.

The ability to exchange A domains with greater accuracy should aid engineering efforts in the future, however many issues remain. As we have observed in nature, there is unlikely to be a single recombination site that will work in every case. Nevertheless, the growing body of genomic data will continue to reveal evolutionary snapshots akin to the wollamide BGC presented here. Over a decade of sequencing has provided us with only a threadbare sample of microbial genomic diversity and, as a result, we are still largely ignorant of the evolutionary processes that govern small molecule biosynthesis. Fortunately, this means there remains wealth of information yet to be gleaned from Nature.

## Methods

For methods and experimental details, please see the supplementary information.

## Acknowledgements

This work was supported by the Biotechnology and Biological Sciences Research Council (BBSRC) via Strategic Program Project BBS/E/J/000PR9790 to the John Innes Centre (JIC), and by Norwich Research Park Doctoral Training Program Studentship BB/M011216/1. We also acknowledge the support of the Cooperative Research Centres Projects Scheme (CRCPFIVE000119). We thank the Earlham Institute (Norwich UK) for sequencing and assembly of the *Streptomyces* sp. MST110588 genome. We also thank the JIC metabolomics platform at for excellent mass spectrometry support.

## Contributions

B.W. and E.L. conceived the study. B.W., T.J.B., K.A.J.B. and J.L. designed the experiments, and T.J.B., K.A.J.B. and J.L. performed the experiments and analysed the data. B.W. and T.J.B. wrote the paper and all authors edited the paper.

## References

(1) Nivina, A.; Yuet, K. P.; Hsu, J.; Khosla, C. Evolution and Diversity of Assembly-Line Polyketide Synthases. Chemical Reviews. American Chemical Society December 26, 2019, pp 12524–12547. https://doi.org/10.1021/acs.chemrev.9b00525.

(2) Süssmuth, R. D.; Mainz, A. Nonribosomal Peptide Synthesis-Principles and Prospects. Angew. Chemie Int. Ed. 2017, 56 (14), 3770–3821. https://doi.org/10.1002/anie.201609079.

(3) Kirst, H. A. The Spinosyn Family of Insecticides: Realizing the Potential of Natural Products Research. Journal of Antibiotics. Nature Publishing Group March 12, 2010, pp 101–111. https://doi.org/10.1038/ja.2010.5.

(4) Fierro, F.; García-Estrada, C.; Castillo, N. I.; Rodríguez, R.; Velasco-Conde, T.; Martín, J. F. Transcriptional and Bioinformatic Analysis of the 56.8 Kb DNA Region Amplified in Tandem Repeats Containing the Penicillin Gene Cluster in Penicillium Chrysogenum. Fungal Genet. Biol. 2006, 43 (9), 618–629. https://doi.org/10.1016/j.fgb.2006.03.001.

(5) Ray, L.; Moore, B. S. Recent Advances in the Biosynthesis of Unusual Polyketide Synthase Substrates. Nat. Prod. Rep. 2016, 33 (2), 150–161. https://doi.org/10.1039/c5np00112a.

(6) Caboche, S.; Leclère, V.; Pupin, M.; Kucherov, G.; Jacques, P. Diversity of Monomers in Nonribosomal Peptides: Towards the Prediction of Origin and Biological Activity. J. Bacteriol. 2010, 192 (19), 5143–5150. https://doi.org/10.1128/JB.00315-10.

(7) Walsh, C. T.; O’Brien, R. V; Khosla, C. Nonproteinogenic Amino Acid Building Blocks for Nonribosomal Peptide and Hybrid Polyketide Scaffolds. Angew. Chem. Int. Ed. Engl. 2013, 52 (28), 7098–7124. https://doi.org/10.1002/anie.201208344.

(8) Blin, K.; Wolf, T.; Chevrette, M. G.; Lu, X.; Schwalen, C. J.; Kautsar, S. A.; Suarez Duran, H. G.; de los Santos, E. L. C., Kim, H. U.; Nave, M.; et al. AntiSMASH 4.0—Improvements in Chemistry Prediction and Gene Cluster Boundary Identification. Nucleic Acids Res. 2017, 45 (W1), W36–W41. https://doi.org/10.1093/nar/gkx319.

(9) Skinnider, M. A.; Merwin, N. J.; Johnston, C. W.; Magarvey, N. A. PRISM 3: Expanded Prediction of Natural Product Chemical Structures from Microbial Genomes. Nucleic Acids Res. 2017, 45 (W1), W49–W54. https://doi.org/10.1093/nar/gkx320.

(10) Dutta, S.; Whicher, J. R.; Hansen, D. A.; Hale, W. A.; Chemler, J. A.; Congdon, G. R.; Narayan, A. R. H.; Håkansson, K.; Sherman, D. H.; Smith, J. L.; et al. Structure of a Modular Polyketide Synthase. Nature 2014, 510 (7506), 512–517. https://doi.org/10.1038/nature13423.

(11) Whicher, J. R.; Dutta, S.; Hansen, D. A.; Hale, W. A.; Chemler, J. A.; Dosey, A. M.; Narayan, A. R. H.; Håkansson, K.; Sherman, D. H.; Smith, J. L.; et al. Structural Rearrangements of a Polyketide Synthase Module during Its Catalytic Cycle. Nature 2014, 510 (7506), 560–564. https://doi.org/10.1038/nature13409.

(12) Drake, E. J.; Miller, B. R.; Shi, C.; Tarrasch, J. T.; Sundlov, J. A.; Leigh Allen, C.; Skiniotis, G.; Aldrich, C. C.; Gulick, A. M. Structures of Two Distinct Conformations of Holo-Non-Ribosomal Peptide Synthetases. Nature 2016, 529 (7585), 235–238. https://doi.org/10.1038/nature16163.

(13) Miller, B. R.; Drake, E. J.; Shi, C.; Aldrich, C. C.; Gulick, A. M. Structures of a Nonribosomal Peptide Synthetase Module Bound to MbtH-like Proteins Support a Highly Dynamic Domain Architecture. J. Biol. Chem. 2016, 291 (43), 22559–22571. https://doi.org/10.1074/jbc.M116.746297.

(14) Bozhüyük, K. A.; Micklefield, J.; Wilkinson, B. Engineering Enzymatic Assembly Lines to Produce New Antibiotics. Current Opinion in Microbiology. Elsevier Ltd October 1, 2019, pp 88–96. https://doi.org/10.1016/j.mib.2019.10.007.

(15) Jenke-Kodama, H.; Dittmann, E. Evolution of Metabolic Diversity: Insights from Microbial Polyketide Synthases. Phytochemistry 2009, 70 (15–16), 1858–1866. https://doi.org/10.1016/J.PHYTOCHEM.2009.05.021.

(16) Zhang, L.; Hashimoto, T.; Qin, B.; Hashimoto, J.; Kozone, I.; Kawahara, T.; Okada, M.; Awakawa, T.; Ito, T.; Asakawa, Y.; et al. Characterization of Giant Modular PKSs Provides Insight into Genetic Mechanism for Structural Diversification of Aminopolyol Polyketides. Angew. Chemie Int. Ed. 2017, 56 (7), 1740–1745. https://doi.org/10.1002/anie.201611371.

(17) Wlodek, A.; Kendrew, S. G.; Coates, N. J.; Hold, A.; Pogwizd, J.; Rudder, S.; Sheehan, L. S.; Higginbotham, S. J.; Stanley-Smith, A. E.; Warneck, T.; et al. Diversity Oriented Biosynthesis via Accelerated Evolution of Modular Gene Clusters. Nat. Commun. 2017, 8 (1), 1206. https://doi.org/10.1038/s41467-017-01344-3.

(18) Baunach, M.; Chowdhury, S.; Stallforth, P.; Dittmann, E. The Landscape of Recombination Events That Create Nonribosomal Peptide Diversity. Mol. Biol. Evol. 2021. https://doi.org/10.1093/molbev/msab015.

(19) Khalil, Z. G.; Salim, A. A.; Lacey, E.; Blumenthal, A.; Capon, R. J. Wollamides: Antimycobacterial Cyclic Hexapeptides from an Australian Soil Streptomyces. Org. Lett. 2014, 16 (19), 5120–5123. https://doi.org/10.1021/ol502472c.

(20) Asfaw, H.; Laqua, K.; Walkowska, A. M.; Cunningham, F.; Martinez-Martinez, M. S.; Cuevas-Zurita, J. C.; Ballell-Pages, L.; Imming, P. Design, Synthesis and Structure-Activity Relationship Study of Wollamide B; a New Potential Anti TB Agent. PLoS One 2017, 12 (4), e0176088. https://doi.org/10.1371/journal.pone.0176088.

(21) Tsutsumi, L. S.; Elmore, J. M.; Dang, U. T.; Wallace, M. J.; Marreddy, R.; Lee, R. B.; Tan, G. T.; Hurdle, J. G.; Lee, R. E.; Sun, D. Solid-Phase Synthesis and Antibacterial Activity of Cyclohexapeptide Wollamide B Analogs. ACS Comb. Sci. 2018, 20 (3), 172–185. https://doi.org/10.1021/acscombsci.7b00189.

(22) Prior, A. M.; Sun, D. Solid-Phase Synthesis of Wollamide Cyclohexapeptide Analogs. In Methods in Molecular Biology; Humana Press Inc., 2020; Vol. 2103, pp 175–187. https://doi.org/10.1007/978-1-0716-0227-0_11.

(23) Song, Y.; Li, Q.; Liu, X.; Chen, Y.; Zhang, Y.; Sun, A.; Zhang, W.; Zhang, J.; Ju, J. Cyclic Hexapeptides from the Deep South China Sea-Derived Streptomyces Scopuliridis SCSIO ZJ46 Active Against Pathogenic Gram-Positive Bacteria. J. Nat. Prod. 2014, 77 (8), 1937–1941. https://doi.org/10.1021/np500399v.

(24) Li, Q.; Song, Y.; Qin, X.; Zhang, X.; Sun, A.; Ju, J. Identification of the Biosynthetic Gene Cluster for the Anti-Infective Desotamides and Production of a New Analogue in a Heterologous Host. J. Nat. Prod. 2015, 78 (4), 944–948. https://doi.org/10.1021/acs.jnatprod.5b00009.

(25) Eid, J.; Fehr, A.; Gray, J.; Luong, K.; Lyle, J.; Otto, G.; Peluso, P.; Rank, D.; Baybayan, P.; Bettman, B.; et al. Real-Time DNA Sequencing from Single Polymerase Molecules. Science 2009, 323 (5910), 133–138. https://doi.org/10.1126/science.1162986.

(26) Chin, C.-S.; Alexander, D. H.; Marks, P.; Klammer, A. A.; Drake, J.; Heiner, C.; Clum, A.; Copeland, A.; Huddleston, J.; Eichler, E. E.; et al. Nonhybrid, Finished Microbial Genome Assemblies from Long-Read SMRT Sequencing Data. Nat. Methods 2013, 10 (6), 563–569. https://doi.org/10.1038/nmeth.2474.

(27) Felnagle, E. A.; Barkei, J. J.; Park, H.; Podevels, A. M.; McMahon, M. D.; Drott, D. W.; Thomas, M. G. MbtH-like Proteins as Integral Components of Bacterial Nonribosomal Peptide Synthetases. Biochemistry 2010, 49 (41), 8815–8817. https://doi.org/10.1021/bi1012854.

(28) Baltz, R. H. Function of MbtH Homologs in Nonribosomal Peptide Biosynthesis and Applications in Secondary Metabolite Discovery. J. Ind. Microbiol. Biotechnol. 2011, 38 (11), 1747–1760. https://doi.org/10.1007/s10295-011-1022-8.

(29) Stachelhaus, T.; Mootz, H. D.; Marahiel, M. A. The Specificity-Conferring Code of Adenylation Domains in Nonribosomal Peptide Synthetases. Chem. Biol. 1999, 6 (8), 493–505. https://doi.org/10.1016/S1074-5521(99)80082-9.

(30) Blin, K.; Shaw, S.; Steinke, K.; Villebro, R.; Ziemert, N.; Lee, S. Y.; Medema, M. H.; Weber, T. AntiSMASH 5.0: Updates to the Secondary Metabolite Genome Mining Pipeline. Nucleic Acids Res. 2019, 47 (W1), W81–W87.

(31) Kouprina, N.; Larionov, V. TAR Cloning: Insights into Gene Function, Long-Range Haplotypes and Genome Structure and Evolution. Nat. Rev. Genet. 2006, 7 (10), 805–812.

(32) Nijkamp, J. F.; van den Broek, M.; Datema, E.; de Kok, S.; Bosman, L.; Luttik, M. A.; Daran-Lapujade, P.; Vongsangnak, W.; Nielsen, J.; Heijne, W. H. M.; et al. De Novo Sequencing, Assembly and Analysis of the Genome of the Laboratory Strain Saccharomyces Cerevisiae CEN.PK113-7D, a Model for Modern Industrial Biotechnology. Microb. Cell Fact. 2012, 11. https://doi.org/10.1186/1475-2859-11-36.

(33) Kieser, T.; Bibb, M. J.; Buttner, M. J.; Chater, K. F.; Hopwood, D. A.; John Innes Foundation. Practical Streptomyces Genetics; John Innes Foundation, 2000.

(34) Hahn, M.; Stachelhaus, T. Harnessing the Potential of Communication-Mediating Domains for the Biocombinatorial Synthesis of Nonribosomal Peptides. Proc. Natl. Acad. Sci. U. S. A. 2006, 103 (2), 275–280. https://doi.org/10.1073/pnas.0508409103.

(35) Hacker, C.; Cai, X.; Kegler, C.; Zhao, L.; Weickhmann, A. K.; Wurm, J. P.; Bode, H. B.; Wöhnert, J. Structure-Based Redesign of Docking Domain Interactions Modulates the Product Spectrum of a Rhabdopeptide-Synthesizing NRPS. Nat. Commun. 2018, 9 (1), 1–11. https://doi.org/10.1038/s41467-018-06712-1.

(36) Watzel, J.; Hacker, C.; Duchardt-Ferner, E.; Bode, H. B.; Wöhnert, J. A New Docking Domain Type in the Peptide-Antimicrobial-Xenorhabdus Peptide Producing Nonribosomal Peptide Synthetase from Xenorhabdus Bovienii. ACS Chem. Biol. 2020. https://doi.org/10.1021/ACSCHEMBIO.9B01022.

(37) Innan, H.; Kondrashov, F. The Evolution of Gene Duplications: Classifying and Distinguishing between Models. Nature Reviews Genetics. Nature Publishing Group February 6, 2010, pp 97–108. https://doi.org/10.1038/nrg2689.

(38) Crüsemann, M.; Kohlhaas, C.; Piel, J. Evolution-Guided Engineering of Nonribosomal Peptide Synthetase Adenylation Domains. Chem. Sci. 2013, 4 (3), 1041–1045. https://doi.org/10.1039/C2SC21722H.

(39) Kries, H.; Niquille, D. L.; Hilvert, D. A Subdomain Swap Strategy for Reengineering Nonribosomal Peptides. Chem. Biol. 2015, 22 (5), 640–648.

(40) Calcott, M. J.; Owen, J. G.; Ackerley, D. F. Efficient Rational Modification of Non-Ribosomal Peptides by Adenylation Domain Substitution. Nat. Commun. 2020, 11 (1). https://doi.org/10.1038/s41467-020-18365-0.

(41) Lynch, M. Streamlining and Simplification of Microbial Genome Architecture. Annu. Rev. Microbiol. 2006, 60 (1), 327–349. https://doi.org/10.1146/annurev.micro.60.080805.142300.

(42) McDonald, B. R.; Currie, C. R. Lateral Gene Transfer Dynamics in the Ancient Bacterial Genus Streptomyces. MBio 2017, 8 (3), e00644–17. https://doi.org/10.1128/mBio.00644-17.

(43) Martin, D. P.; Murrell, B.; Golden, M.; Khoosal, A.; Muhire, B. RDP4: Detection and Analysis of Recombination Patterns in Virus Genomes. Virus Evol. 2015, 1 (1), vev003. https://doi.org/10.1093/ve/vev003.

(44) Medema, M. H.; Kottmann, R.; Yilmaz, P.; Cummings, M.; Biggins, J. B.; Blin, K.; De Bruijn, I.; Chooi, Y. H.; Claesen, J.; Coates, R. C.; et al. Minimum Information about a Biosynthetic Gene Cluster. Nature Chemical Biology. Nature Publishing Group August 18, 2015, pp 625–631. https://doi.org/10.1038/nchembio.1890.

(45) Gilchrist, C. L. M.; Booth, T. J.; Chooi, Y.-H. Cblaster: A Remote Search Tool for Rapid Identification and Visualisation of Homologous Gene Clusters. bioRxiv 2020, 2020.11.08.370601. https://doi.org/10.1101/2020.11.08.370601.

(46) McClure, R. A.; Goering, A. W.; Ju, K. S.; Baccile, J. A.; Schroeder, F. C.; Metcalf, W. W.; Thomson, R. J.; Kelleher, N. L. Elucidating the Rimosamide-Detoxin Natural Product Families and Their Biosynthesis Using Metabolite/Gene Cluster Correlations. ACS Chem. Biol. 2016, 11 (12), 3452–3460. https://doi.org/10.1021/acschembio.6b00779.

(47) Kadi, N.; Challis, G. L. Chapter 17 Siderophore Biosynthesis. In Methods in enzymology; 2009; Vol. 458, pp 431–457. https://doi.org/10.1016/S0076-6879(09)04817-4.

(48) Jenke-Kodama, H.; Sandmann, A.; Müller, R.; Dittmann, E. Evolutionary Implications of Bacterial Polyketide Synthases. Mol. Biol. Evol. 2005, 22 (10), 2027–2039. https://doi.org/10.1093/molbev/msi193.

(49) Cimermancic, P.; Medema, M. H.; Claesen, J.; Kurita, K.; Wieland Brown, L. C., Mavrommatis, K.; Pati, A.; Godfrey, P. A.; Koehrsen, M.; Clardy, J.; et al. Insights into Secondary Metabolism from a Global Analysis of Prokaryotic Biosynthetic Gene Clusters. Cell 2014, 158 (2), 412–421. https://doi.org/10.1016/j.cell.2014.06.034.

(50) Zhou, Y.; Lin, X.; Xu, C.; Shen, Y.; Wang, S. P.; Liao, H.; Li, L.; Deng, H.; Lin, H. W. Investigation of Penicillin Binding Protein (PBP)-like Peptide Cyclase and Hydrolase in Surugamide Non-Ribosomal Peptide Biosynthesis. Cell Chem. Biol. 2019, 26 (5), 737- 744.e4. https://doi.org/10.1016/j.chembiol.2019.02.010.

(51) Fazal, A.; Webb, M. E.; Seipke, R. F. The Desotamide Family of Antibiotics. Antibiotics. MDPI AG August 1, 2020, pp 1–14. https://doi.org/10.3390/antibiotics9080452.

(52) Conrad, B.; Antonarakis, S. E. Gene Duplication: A Drive for Phenotypic Diversity and Cause of Human Disease. 2007. https://doi.org/10.1146/annurev.genom.8.021307.110233.

(53) Qian, W.; Zhang, J. Genomic Evidence for Adaptation by Gene Duplication. Genome Res. 2014, 24 (8), 1356–1362. https://doi.org/10.1101/gr.172098.114.

(54) Medema, M. H.; Cimermancic, P.; Sali, A.; Takano, E.; Fischbach, M. A. A Systematic Computational Analysis of Biosynthetic Gene Cluster Evolution: Lessons for Engineering Biosynthesis. PLoS Comput. Biol. 2014, 10 (12), e1004016. https://doi.org/10.1371/journal.pcbi.1004016.

(55) Román-Hurtado, F.; Sánchez-Hidalgo, M.; Martín, J.; Javier Ortiz-López, F.; Reyes, F.; Genilloud, O. One Pathway, Two Cyclic Pentapeptides: Heterologous Expression of BE-18257 A-C and 1 Pentaminomycins A-E from Streptomyces Cacaoi CA-170360 2 3. bioRxiv 2020, 2020.10.23.352575. https://doi.org/10.1101/2020.10.23.352575.

(56) Liao, D. Concerted Evolution: Molecular Mechanism and Biological Implications. Am. J. Hum. Genet. 1999, 64 (1), 24–30. https://doi.org/10.1086/302221.

(57) Santoyo, G.; Romero, D. Gene Conversion and Concerted Evolution in Bacterial Genomes. FEMS Microbiol. Rev. 2005, 29 (2), 169–183. https://doi.org/10.1016/j.femsre.2004.10.004.

(58) Niquille, D. L.; Hansen, D. A.; Mori, T.; Fercher, D.; Kries, H.; Hilvert, D. Nonribosomal Biosynthesis of Backbone-Modified Peptides. Nat. Chem. 2018, 10 (3), 282–287. https://doi.org/10.1038/nchem.2891.

(59) Bozhüyük, K. A. J.; Fleischhacker, F.; Linck, A.; Wesche, F.; Tietze, A.; Niesert, C.-P.; Bode, H. B. De Novo Design and Engineering of Non-Ribosomal Peptide Synthetases. Nat. Chem. 2017. https://doi.org/10.1038/nchem.2890.

(60) Bozhüyük, K. A. J.; Linck, A.; Tietze, A.; Kranz, J.; Wesche, F.; Nowak, S.; Fleischhacker, F.; Shi, Y. N.; Grün, P.; Bode, H. B. Modification and de Novo Design of Non-Ribosomal Peptide Synthetases Using Specific Assembly Points within Condensation Domains. Nat. Chem. 2019, 11 (7), 653–661. https://doi.org/10.1038/s41557-019-0276-z.

(61) Bozhueyuek, K. A. J.; Watzel, J.; Abbood, N.; Bode, H. B. Synthetic Zippers as an Enabling Tool for Engineering of Non-Ribosomal Peptide Synthetases. bioRxiv. bioRxiv May 6, 2020, p 2020.05.06.080655. https://doi.org/10.1101/2020.05.06.080655.

(62) Atanasov, A. G.; Zotchev, S. B.; Dirsch, V. M.; Orhan, I. E.; Banach, M.; Rollinger, J. M.; Barreca, D.; Weckwerth, W.; Bauer, R.; Bayer, E. A.; et al. Natural Products in Drug Discovery: Advances and Opportunities. Nature Reviews Drug Discovery. Nature Research March 1, 2021, pp 200–216. https://doi.org/10.1038/s41573-020-00114-z.

(63) Meyer, S.; Kehr, J. C.; Mainz, A.; Dehm, D.; Petras, D.; Süssmuth, R. D.; Dittmann, E. Biochemical Dissection of the Natural Diversification of Microcystin Provides Lessons for Synthetic Biology of NRPS. Cell Chem. Biol. 2016, 23 (4), 462–471. https://doi.org/10.1016/j.chembiol.2016.03.011.

(64) Scaglione, A.; Fullone, M. R.; Montemiglio, L. C.; Parisi, G.; Zamparelli, C.; Vallone, B.; Savino, C.; Grgurina, I. Structure of the Adenylation Domain Thr1 Involved in the Biosynthesis of 4-Chlorothreonine in Streptomyces Sp. OH-5093—Protein Flexibility and Molecular Bases of Substrate Specificity. FEBS J. 2017, 284 (18), 2981–2999. https://doi.org/10.1111/febs.14163.

(65) Sundlov, J. A.; Shi, C.; Wilson, D. J.; Aldrich, C. C.; Gulick, A. M. Structural and Functional Investigation of the Intermolecular Interaction between NRPS Adenylation and Carrier Protein Domains. Chem. Biol. 2012, 19 (2), 188–198. https://doi.org/10.1016/j.chembiol.2011.11.013.

(66) Tanovic, A.; Samel, S. A.; Essen, L.-O.; Marahiel, M. A. Crystal Structure of the Termination Module of a Nonribosomal Peptide Synthetase. Science 2008, 321 (5889), 659–663. https://doi.org/10.1126/science.1159850.

(67) Reimer, J. M.; Haque, A. S.; Tarry, M. J.; Schmeing, T. M. Piecing Together Nonribosomal Peptide Synthesis. Curr. Opin. Struct. Biol. 2018, 49, 104–113. https://doi.org/10.1016/J.SBI.2018.01.011.

(68) Reimer, J. M.; Eivaskhani, M.; Harb, I.; Guarné, A.; Weigt, M.; Martin Schmeing; T. Structures of a Dimodular Nonribosomal Peptide Synthetase Reveal Conformational Flexibility. Science (80-.). 2019, 366 (6466). https://doi.org/10.1126/science.aaw4388.

(69) Lautru, S.; Challis, G. L. Substrate Recognition by Nonribosomal Peptide Synthetase Multi-Enzymes. Microbiology. Society for General Microbiology June 1, 2004, pp 1629–1636. https://doi.org/10.1099/mic.0.26837-0.

(70) Baltz, R. H. Combinatorial Biosynthesis of Cyclic Lipopeptide Antibiotics: A Model for Synthetic Biology to Accelerate the Evolution of Secondary Metabolite Biosynthetic Pathways. ACS Synthetic Biology. American Chemical Society October 17, 2014, pp 748–758. https://doi.org/10.1021/sb3000673.

(71) Belshaw, P. J.; Walsh, C. T.; Stachelhaus, T. Aminoacyl-CoAs as Probes of Condensation Domain Selectivity in Nonribosomal Peptide Synthesis. Science. 1999, 284 (5413), 486–489. https://doi.org/10.1126/science.284.5413.486.

(72) Linne, U.; Marahiel, M. A. Control of Directionality in Nonribosomal Peptide Synthesis: Role of the Condensation Domain in Preventing Misinitiation and Timing of Epimerization. Biochemistry 2000, 39 (34), 10439–10447. https://doi.org/10.1021/bi000768w.

(73) Fewer, D. P.; Rouhiainen, L.; Jokela, J.; Wahlsten, M.; Laakso, K.; Wang, H.; Sivonen, K. Recurrent Adenylation Domain Replacement in the Microcystin Synthetase Gene Cluster. BMC Evol. Biol. 2007, 7 (1), 183. https://doi.org/10.1186/1471-2148-7-183.

(74) Conti, E.; Stachelhaus, T.; Marahiel, M. A.; Brick, P. Structural Basis for the Activation of Phenylalanine in the Non-Ribosomal Biosynthesis of Gramicidin S. EMBO J. 1997, 16 (14), 4174–4183.

