## Supplementary information for "Bifurcation drives the evolution of assembly-line biosynthesis"

Thomas J. Booth, Kenan A. J. Bozhüyük, Jonathon D. Liston, Ernest Lacey & Barrie Wilkinson*

### 1. Supplementary Methods

#### 1.1 Strains and culture conditions

All strains and plasmids used in the study can be found in Supplementary Tables 12 and 13. *E. coli* strains were cultured on LB medium (25 g/L LB broth (Miller)) at 37 °C. *S. cerevisae* CEN.PK 2-1c and derivatives were cultured on YPDA (10 g/L yeast extract, 20 g/L peptone and 20 g/L glucose) supplemented with 0.004 % adenine sulfate. *Streptomyces* spp. were cultured on SF+M agar (20 g/L soya flour, 20 g/L mannitol, 20 g/L Lab M No 1 agar) at 30 °C or in SV2 (glucose 15 g/L, glycerol 15 g/L, soy peptone 15 g/L, sodium chloride 3 g/L, calcium carbonate 1 g/L). Apramycin (100 µg/ml), kanamycin (50 µg/ml) and G418 (200 µg/ml) were used as selection markers.

MST-70754, MST-71321, MST-71458, MST-94754 and MST-127221 were isolated by Microbial Screening Technologies (Smithsfield, NSW, Australia) from soil samples collected from across New South Wales between 1994 and 1998.

#### 1.2 Genome Sequencing and Genomic Analysis

High molecular weight genomic DNA was extracted according to a modified version the salting out procedure described by Kieser et al.[1] with the modifications as described here: wet mycelium (0.5 mL) from a 30 h old SV2 culture was washed with 10% sucrose (10 mL) before resuspension in SET buffer (5 mL; 75 mM NaCl, 25 mM EDTA, 20 mM TrisHCl pH 8.0) to which lysozyme (200 µL; 50 mg/mL) and ribonuclease A (15 µL; 10 mg/mL) were added. The cells were incubated overnight at 37°C; fresh lysozyme (300 µL) was added after ca. 17 h followed by an additional 2 h incubation.

Genomic DNA of *Streptomyces* sp. MST-110588 was sequenced with Pacific Biosciences (PacBio) RSII SMRT technology (commissioned to the Earlham Institute Norwich, UK) and assembled *via* the HGAP2.0 pipeline. Genomic DNA of *Streptomyces* spp. MST-70754, MST-71321, MST-71458, MST-94754 and MST-127221 were sequenced with Illumina MiSeq using paired end sequencing and Nextera Mate Pair library preparation and assembled in house (commissioned to the University of Cambridge DNA sequencing facility). The genome sequence of MST-110588 was deposited in GenBank (BioProject: PRJNA727408, accession: CP074380). The desotamide BGCs from MST-70754 (GenBank: MZ093610), MST-71321 (GenBank: MZ093611), MST-71458 (GenBank: MZ093612), MST-94754 (GenBank: MZ093613) and MST-127221 (GenBank: MZ093614) were deposited in GenBank under the respective accessions.

Genomic DNA sequences were annotated using prodigal[2] as implemented by antiSMASH 5.0[3]. Strain taxonomy was performed using multi-locus sequence typing was implemented in AutoMLST[4]. Adenylation domain specificities were predicted by NRPSsp[5] or by NRPSpredictor2[6] and the specificity codes described by Stachelhaus[7] and Minowa[8], as implemented in antiSMASH. Gene homologues were identified from translated protein sequences using BLASTp[9]. Models of the assembly line modules and adenylating domains were generated *via* remote homology detection using Phyre2[10] and visualised using PyMOL[11].

**1.3 Metabolic Analysis**

*Streptomyces* spp. were grown on SF+M agar for 10 days. 1 $cm^2$ agar plugs were taken from the plates and extracted by shaking with methanol (1 ml) for 30 min. Samples were centrifuged at max speed and the supernatant was dried *in vacuo* prior to resuspension in 100 µl methanol. LCMS analysis was performed using a Shimadzu platform equipped with a NexeraX2 liquid chromatograph (LC30AD), a Prominence photo diode array detector (SPD-M20A) and an LCMS-IT-TOF mass spectrometer. Chromatography was achieved over a Kinetex C18 100A column (100 × 2.1 mm, 2.6 µm; Phenomenex) using a water-methanol gradient (20-100% methanol with 0.1 % formic acid over 12 min with 1 min hold at 100%; flow rate, 0.5 ml/min).

**1.4 Evolutionary Analysis of NRPS Assembly-lines**

Condensation domain and adenylation domain coding nucleotide sequences of *Streptomyces* sp. MST-110588, annotated by antiSMASH (see section 1.2), were aligned using ClustalW (cost matrix: BLOSUM, gap open cost: 10, gap extend cost: 0.1)[12]. Recombination detection was performed on the resulting adenylation domain alignments using RDP[13], GENECONV[14], BOOTSCAN[15], MAXCHI[16], CHIMERA[17], SISCAN[18], LARD[19] and 3SEQ[20] as implemented in RDP4[21] using default settings. Breakpoints were plotted on translated protein sequences by manual annotation. Phylogenies were generated using FastTree 2[22].

**1.5 Assembly of the plasmids pBO1 and pBO1-*wolG2***

Plasmid isolation from *E. coli* and *S. cerevisiae* was performed using PureYield Plasmid Miniprep System (Promega) and Zymoprep Yeast Plasmid Miniprep II (Zymo Research) respectively. The plasmid pBO1 is a selectable yeast centromeric-*E. coli-Streptomyces* shuttle plasmid. It is a derivative of pGP9[23]. The yeast cassette from pFF62A (KanMX4 and 2micron2 Ori)[24] was introduced into the pGP9 backbone to produce pBO1. pGP9 was linearized by PCR amplification with Q5 High-Fidelity

DNA polymerase (NEB) using primers pBO1-pGP9-F1 and pBO1-pGP9-R1, and the yeast cassette from pFF62A was amplified with primers pBO1-pFF-F1 and pBO1-pFF-R1 (Supplementary Table 14). The PCR linearized pGP9 and yeast cassette were transformed into *S. cerevisae* CEN.PK 2-1c according to Schiestl and Gietz[25,26]. Plasmid pBO1-*wolG2* was generated by transformation associated homologous recombination (TAR). pBO1 was linearized with NdeI. *wolG2* was amplified with primers pBO1_wolG2_F1 and pBO1_wolG2_R1 (Supplementary Table 14) from gDNA of *Streptomyces* sp. MST-110588). Linear pBO1 and PCR amplified *wolG2* fragments were assembled by transformation into *S. cerevisae* CEN.PK 2-1c. Plasmids were purified from G418 resistant colonies.

##### 1.6 Heterologous expression of *wolG2* and metabolic analysis

Plasmids pBO1 and pBO1-*wolG2* were extracted from *E. coli* DH5α using Wizard Plus SV miniprep system (Promega) and transformed into electrocompetent *E. coli* ET12567/pUZ8002[27]. The resulting strains were used for the conjugation. Conjugation of *Streptomyces* sp. MST-70754 spores was carried out according to Kieser et al.[1] The resulting exconjugants were cultured on SF+M agar plates containing apramycin (100 µg/ml) and extracted with methanol and analysed by LCMS as described above.

##### 1.7 Modelling of Docking Domain Complexes

Protein sequences of WolG1_NDD–WolH_CDD and WolG2_NDD–WolH_CDD as well as the crystal structure coordinates of PaxC_NDD–PaxB_CDD (PDB-ID: 6TRP_1) [28] were loaded into the Molecular Operating Environment (MOE) 2019.0102[29]. Prior to homology modeling the crystal structure of 6TRP_1 was prepared (*i.e.*, wrong protonation, chirality, and hybridization) and a structural alignment was made. A series of 10 models per protein were constructed with MOE using a Boltzmann-weighted randomized procedure combined with specialized logic for the handling of sequence insertions and deletions[30,31]. The model with the best packing quality function was selected for full energy minimization. Using the AMBER14 forcefield parameters for proteins (Amber14: EHT), the calculated MOE packing scores for models of WolG1_NDD–WolH_CDD and WolG2_NDD–WolH_CDD were 2.4286 and 2.2322, respectively. The stereochemical qualities of the models were assessed using Ramachandran plots and by calculating the Root-Mean-Square-Deviation (RMSD) values of the superposed Cα-atoms of the model with its respective template structure. RMSDs of WolG1_NDD–WolH_CDD and WolG2_NDD–WolH_CDD with 6TRP_1 were 0.1724 and 0.1762 respectively.

##### 1.8 Protein expression, purification and biochemical assays

Adenylation domains were assembled into pET28a using the methodology described in section 1.5 using the primers from Supplementary Table 14. Additionally, the MbtH-like protein coding sequence *wolF2* was cloned into pCDF-Duet-1 using the same procedure. *E. coli* 2(DE3)pLysS containing the both expression plasmids were grown overnight in 10 ml LB. Overnight cultures were used to inoculate 1 L LB and grown to OD600 ~0.5, prior to induction with IPTG (1mM). Following induction, cultures were grown overnight at 18°C. Cultures were centrifuged at 9000 RPM for 20 minutes and the supernatant was discarded. Pellets were resuspended in 15-20 ml P-Buffer (1M $K_2HPO_4$ 9.4 %; 1M $KH_2PO_4$, 0.6 %, pH 8.0). Cells were homogenised using an Avestin EmulsiFlex-B15 (40 bar) and the supernatant was removed and filtered.

Proteins were purified on an ÄKTA Pure chromatography system (GE Healthcare) using a His-Trap 1 ml column. The column was equilibrated with P-buffer until a stable UV signal was established. The equilibrated column was sequentially washed with 5 column volumes of P-buffer containing 10 mM, 20 mM and 50 mM imidazole. Proteins were eluted with 24 column volumes of P-buffer containing of 250 mM imidazole. Fractions with an UV absorbance at 280 nm were pooled. The presence of the correct MW protein was assessed by SDS-PAGE using 12 % RunBlue SDS protein gels (Expedeon). Protein concentration was determined by Bradford assay[32] (BIORAD).

Hydroxylamine trapping assays were performed using the method described by Kadi and Challis[33] with minor modifications. Enzyme (50 mM; 30 µl) was added to the assay solution for each amino acid (15 mM $MgCl_2$, 2.25 mM ATP, 150 mM hydroxylamine, 3 mM amino acid). Samples were incubated for 2 h prior to the addition of stopping solution (10 % (w/v) $FeCl_3 \cdot 6H_2O$, 3.3% trichloroacetic acid in 0.7 M HCl). The absorbance at540 nm was measured using a BioMate 3 spectrophotometer (Thermospectronic).

#### 2. Supplementary Figures

**2.1 Supplementary Figure 1:** Cluster Maps of Hexapeptide producing BGCs

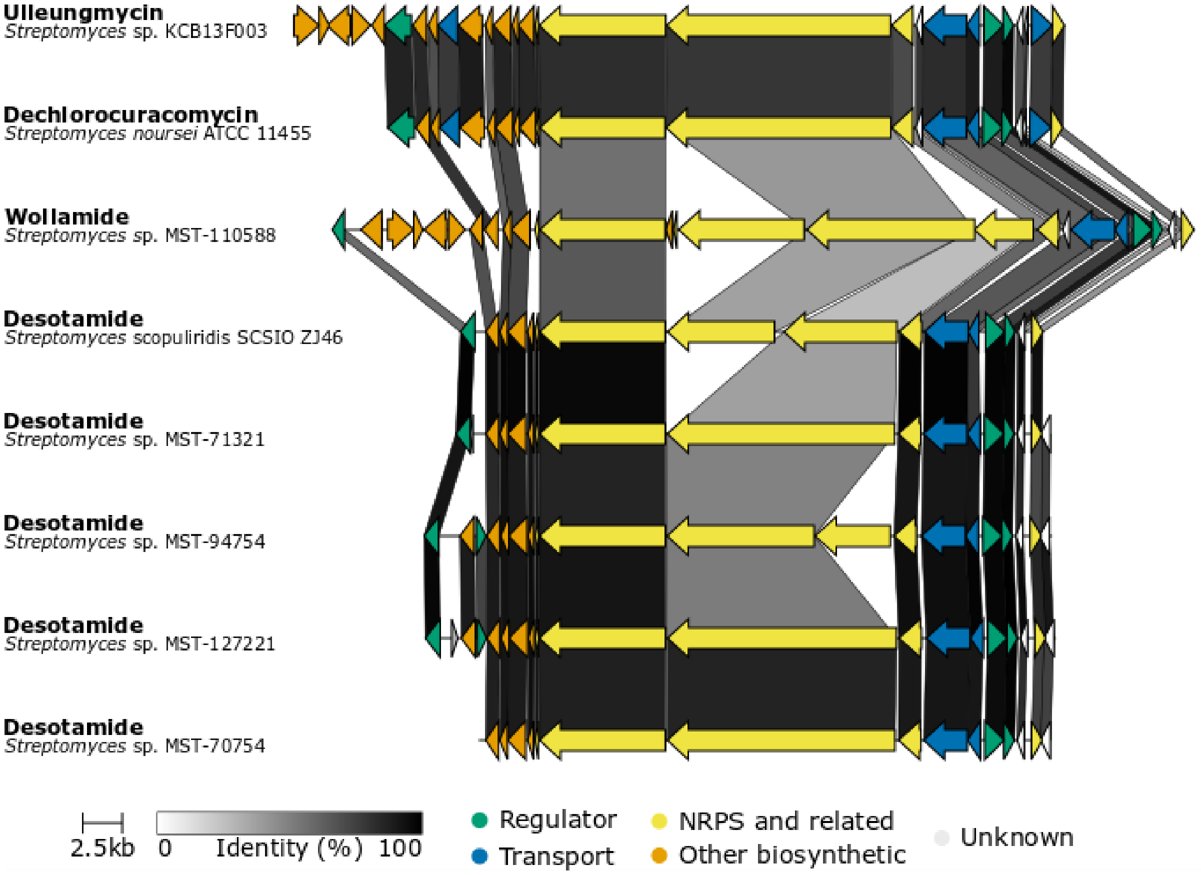

Comparison of hexapeptide producing strains from the Microbial Screening Technologies collection (MST-110588, MST-71321, MST-94754, MST-127221 and MST-70754 (this study)) and previously published hexapeptide BGCs for desotamide[34], dechlorocuracomycin[35] and ulleungmycin[36]. Figure adapted from clinker output[37].

**2.2 Supplementary Figure 2:** Plasmid Map of pBO1.

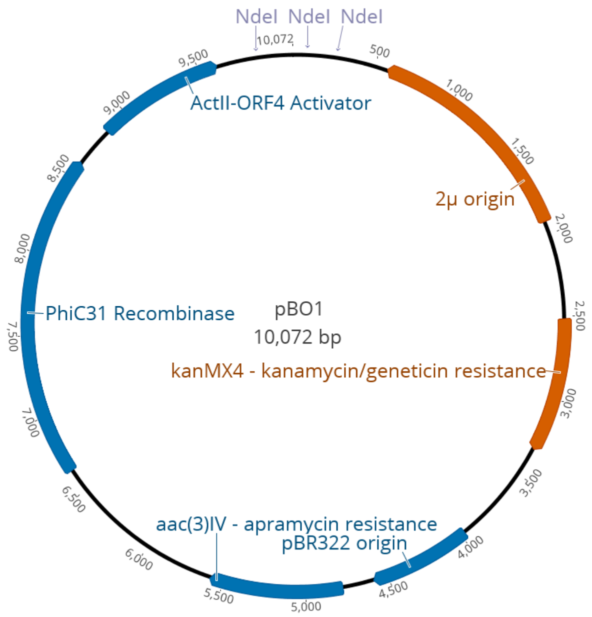

Regions derived from pGP9[23] are coloured blue; those derived from pFF62A[24] are coloured red.

**2.3 Supplementary Figure 3:** LCMS analysis of wollamide and desotamide congeners produced by *Streptomyces* sp. MST-110588

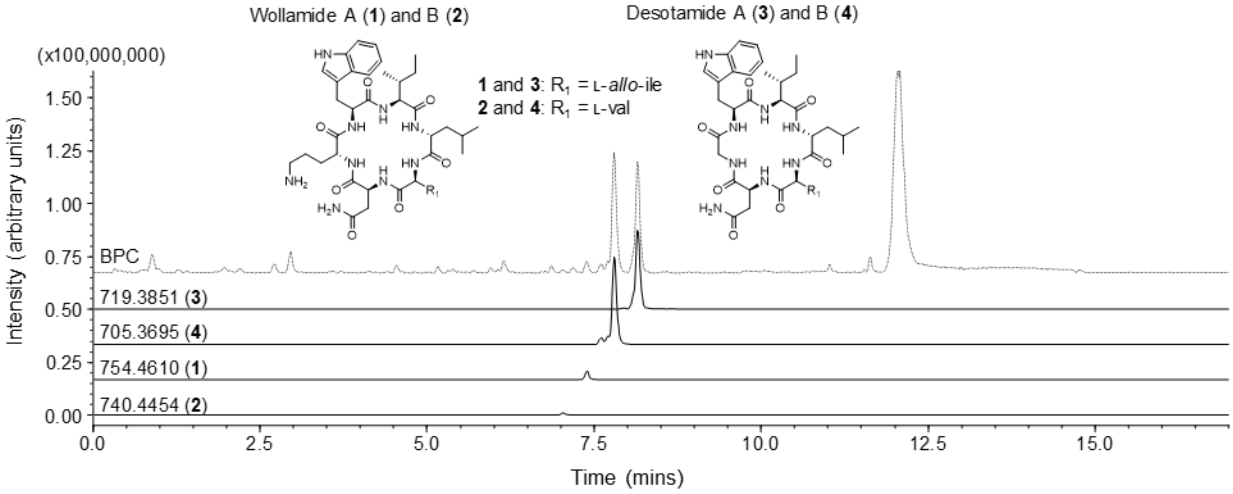

LCMS chromatograms of methanolic fermentation extracts from *Streptomyces* sp. MST-110588. Showing the base peak chromatogram (BPC) plus extracted ion chromatograms for the masses of wollamide A (**1**: $[M+H]^+$ for $C_{38}H_{60}N_9O_7$ = 754.4610) and wollamide B (**2**: $[M+H]^+$ for $C_{37}H_{58}N_9O_7$ = 740.4454), and desotamide A (**3**: exact mass: $[M+Na]^+$ for $C_{35}H_{52}N_8O_7Na$ = 719.3851) and desotamide B (**4**: exact mass: $[M+Na]^+$ for $C_{34}H_{50}N_8O_7Na$ = 705.3695) as previously reported[38].

**2.4 Supplementary Figure 4:** Sequences of docking-domain complexes used for homology modeling.

**6TRP_1|Chain**

MNINEQTLDKLRQAVLQKKIKERIQNSLSTEKY **GSGSGSGSGSGSGSGSGSGSGSGSGY** QIETFFAQDIESVQKELENLSEEELLAMLNGDQQ 93

**WolG1_NDD – linker - WolH_CDD**

MDAAAAARESWIQSLP**E**HVRDRMRSRLTGGAQDG **GSGSGSGSGSGSGSGSGSGSGSGSGY** EIFMIDLPA**R**TLFDNPTVADLAAAVEAQVLAELS 93

**WolG2_NDD -linker - WolH_CDD**

MDAAASRESWIQSLP**A**HVRDRMRSRLAGGAEAG **GSGSGSGSGSGSGSGSGSGSGSGSGY** EIFMIDLPA**R**TLFDNPTVADLAAAVEAQVLAELS 93

Specific interactions between different proteins in NRPS systems are mediated by short C- and N-terminal docking domains ($^{C/N}$DDs). The solution structure 6TRP_1 (PaxC_NDD–PaxB_CDD) was used as a template[39] to model both WolG1_NDD–WolH_CDD and WolG2_NDD–WolH_CDD. As artificial linking of the PaxB/PaxC DD pair via a flexible glycine-serine (GS) linker led to the elucidation of the structure of the DD complex by NMR spectroscopy, we applied an analogous in silico approach for homology modeling. Left: N-terminal DD; Center: GS linker; Right: C-terminal DD. Proposed key interaction of the WolG1 WolH DD pair (red) that is missing in the WolG2 WolH pair (blue) is highlighted (c.f. Supplementary Figure S5).

**2.5 Supplementary Figure 5:** Homology models of WolH/WolG docking-domain complexes.

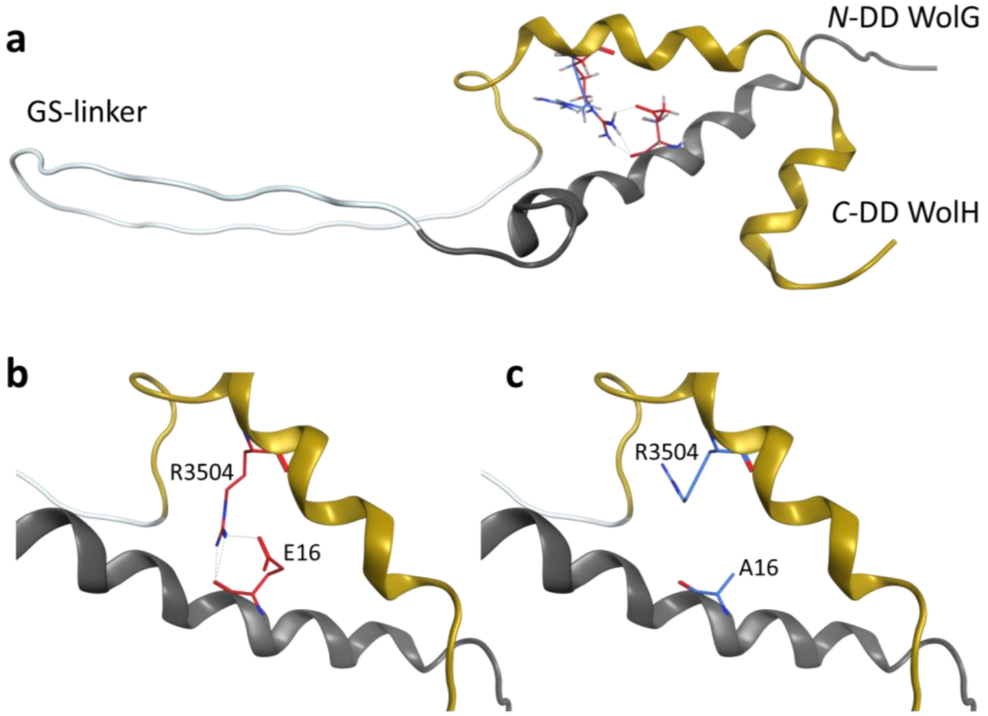

Homology models were calculated to reveal the reason for observed differences in desotamide and wollamide peptide titers. The WolG1 & WolG2 DDs are colored grey, the WolH DD is colored yellow, and the artificial GS linker is colored white. Depicted amino acid residues (stick representation) are colored as introduced in Supplementary Figure S4. (a) Superimposed models of WolG1 WolH and WolG2 WolH DD pairs have an RMSD of 0.051. (b) Excised WolG1 WolH DD pair. Amino acid residues E16 (WolH1) and R3504 (WolH) are forming a salt bridge. (c) Excised WolG2 WolH DD pair. The E16A amino acid change in WolG2 likely causes decreased DD pair affinities. This in turn may serve as an explanation for lower wollamide titers compared to desotamide titers.

**2.6 Supplementary Figure 6:** Pairwise identity of WolG1 and WolG2

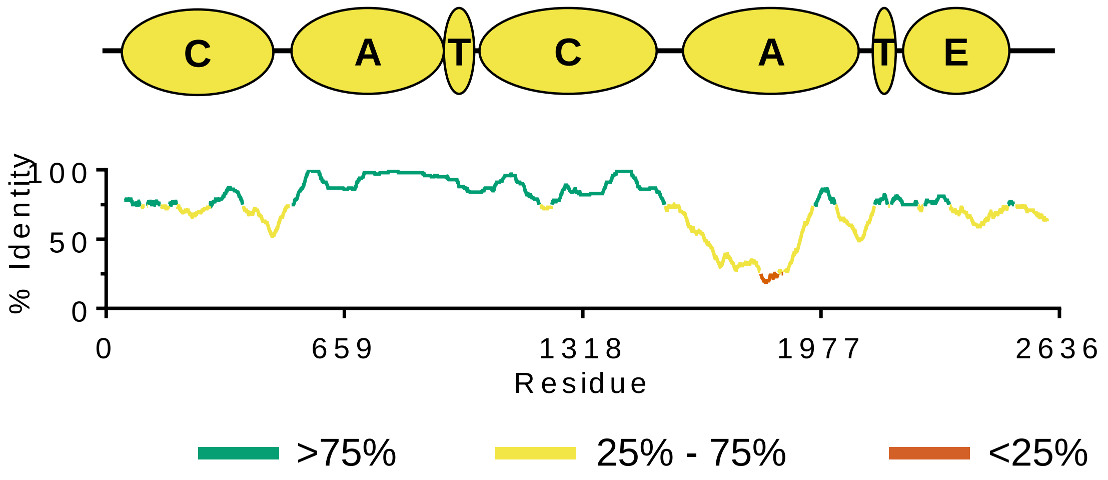

The domain architecture of WolG1 is shown across the top of the figure with the domains denoted as follows: condensation (C), adenylation (A), thiolation/peptidyl carrier protein (T) and, epimerase (E). The identity is also colour coded for convenience. Regions of high homology (>75%) are coloured in green, medium homology in yellow (25 -75%) and low homology in red (<25%).

**2.7 Supplementary Figure 7:** Phylogeny of *Streptomyces* sp. MST-110588 Condensation domains

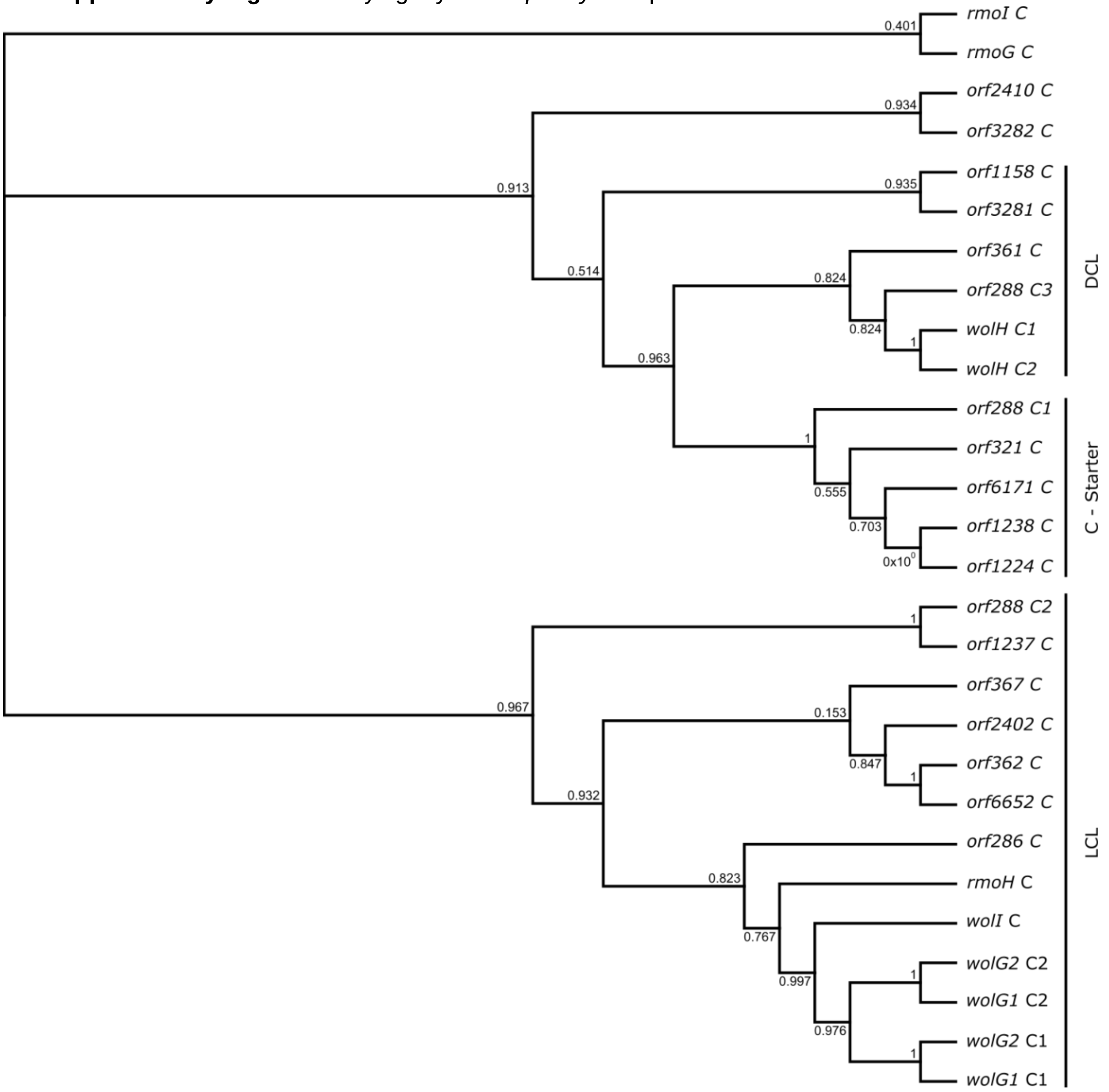

Phylogenetic tree of MST-110588 condensation domains generated by maximum likelihood as implemented in FastTree2[22] from a ClustalW[12] alignment of nucleotide sequences. Condensation domain functions were predicted by NaPDoS[40] major clades are indicated ($^{L}C_{L}$s, which condense two L-amino acid residues; $^{D}C_{L}$s that condense a D-amino acid residues with L-amino acid residues and; C Starter domains, associated with the loading modules of NRPSs).

**2.8 Supplementary Figure 8:** Phylogeny of *Streptomyces* sp. Adenylation domains (recombinant region)

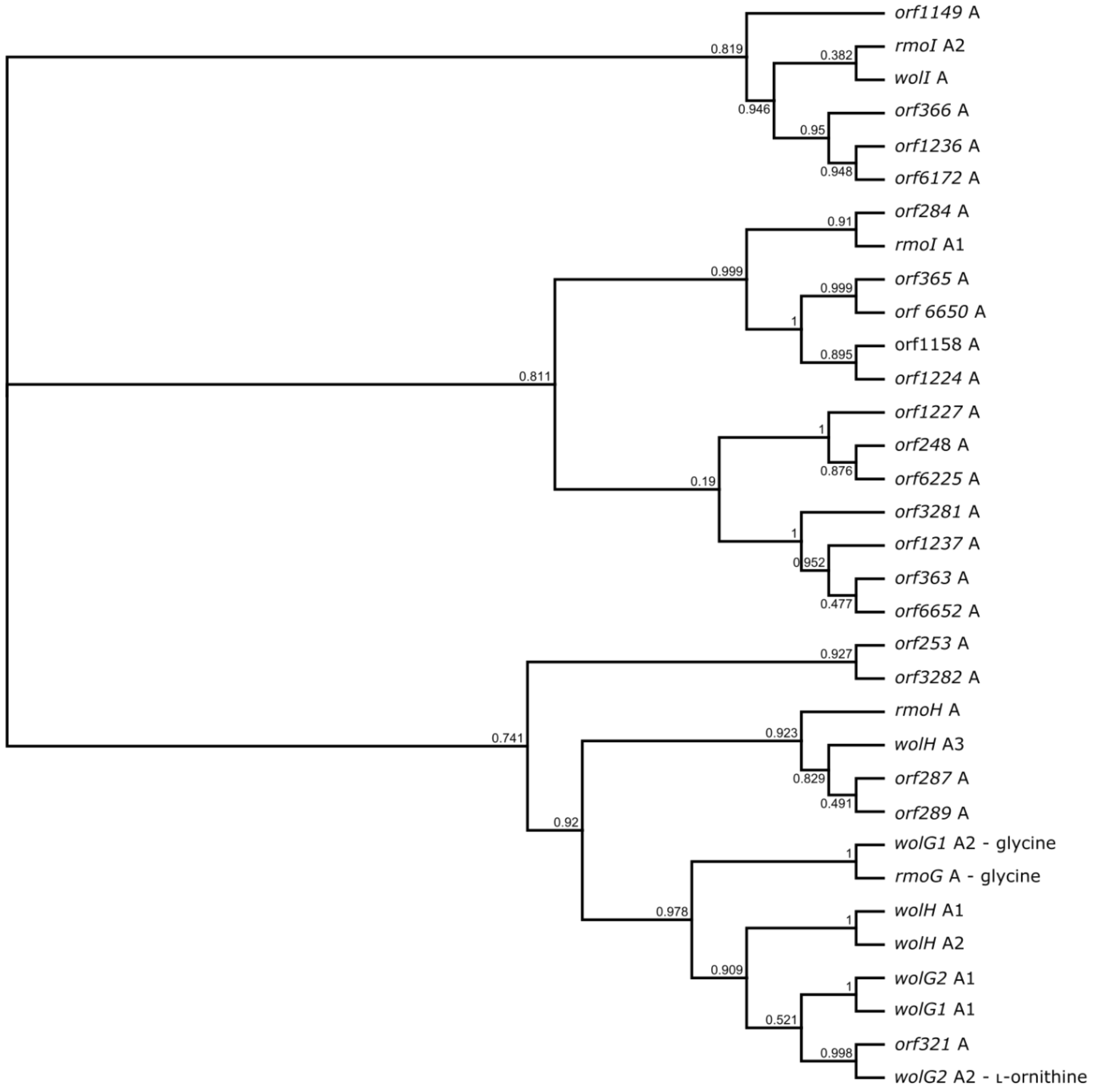

Phylogenetic tree of the *wolG1*/*wolG2/RmoG* recombinant region of MST-110588 adenylation domains (as predicted by RDP4) generated by maximum likelihood as implemented in FastTree2[22] from a ClustalW[12] alignment of nucleotide sequences. Observed substrate specificities encoded by *wolG1*, *wolG2* and *rmoG* are indicated.

**2.9 Supplementary Figure 9:** Phylogeny of *Streptomyces* sp. Adenylation domains (parental region)

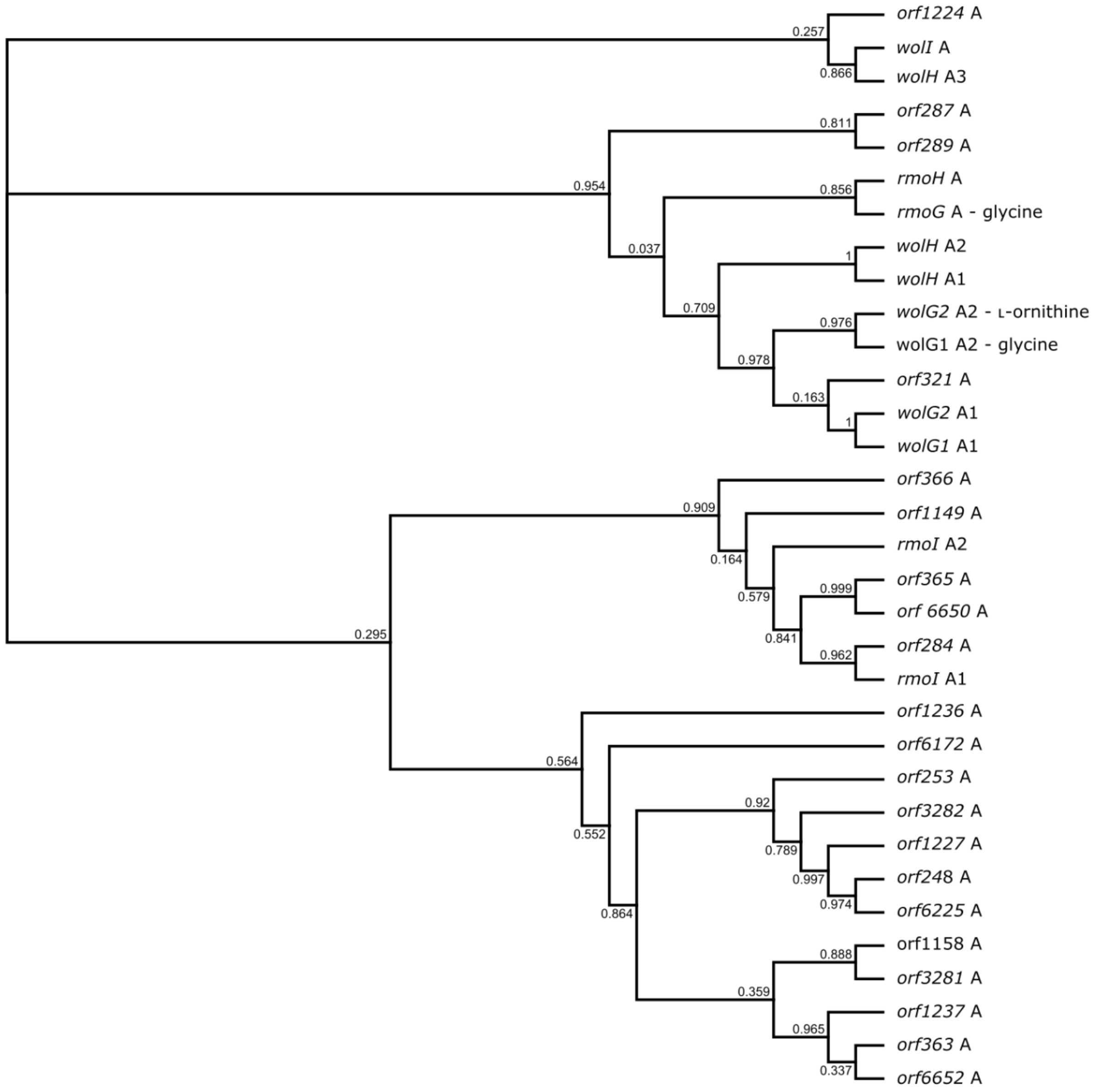

Phylogenetic tree of the of MST-110588 adenylation domains with the *wolG1*/*wolG2/rmoG* recombinant region removed, generated by maximum likelihood as implemented in FastTree2[22] from a ClustalW[12] alignment of nucleotide sequences. Observed substrate specificities encoded by *wolG1*, *wolG2* and *rmoG* are indicated.

**2.10 Supplementary Figure 10:** Topology of a typical adenylation domain showing predicted recombination breakpoints

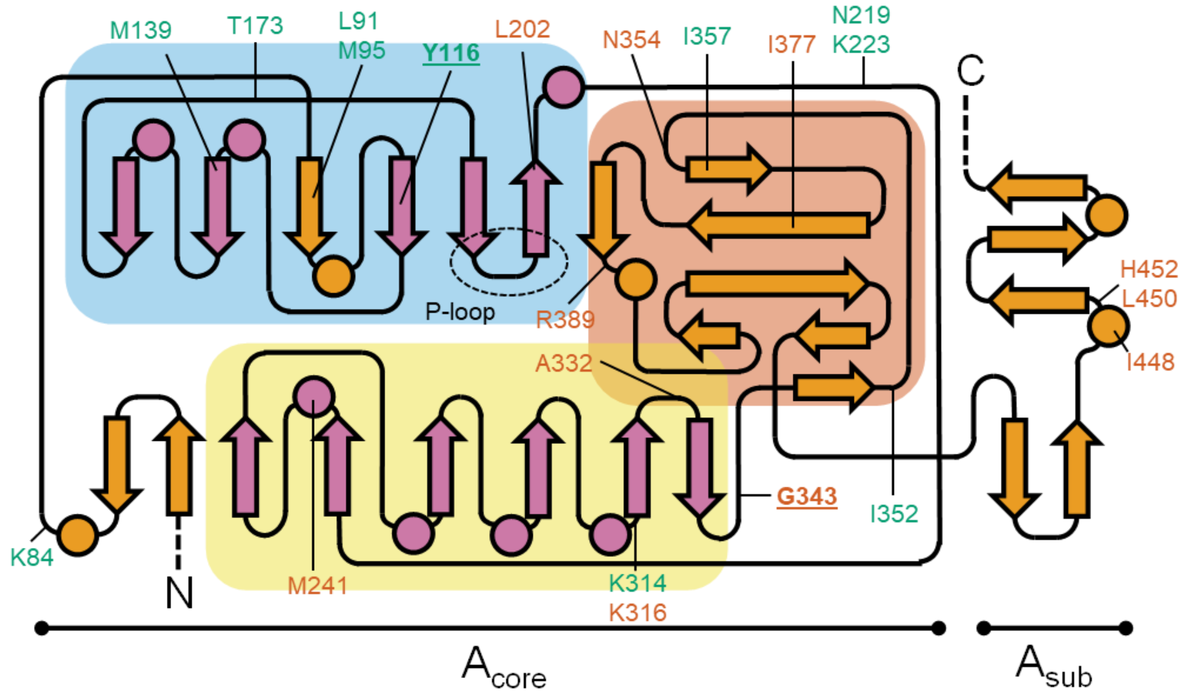

Topology of a typical adenylation domain adapted from Conti et al.[41]. Circles represent α-helices, arrows represent β-strands, and lines represent loops. Regions corresponding to the large *N*-terminal domain ($A_{core}$) and small *C*-terminal domain ($A_{sub}$) are highlighted and subdomains shaded (subdomain 1 = blue; subdomain 2 (flavodoxin-like subdomain) = yellow; subdomain 3 = red). The residues indicated correspond to approximate sites of recombination from Supplementary Table 9 (green = N-terminal; red = C-terminal). Recombinant region of *wolG1* coloured in purple and associated breakpoints are underlined.

**2.11 Supplementary Figure 11:** Homology model of the final module of WolG1

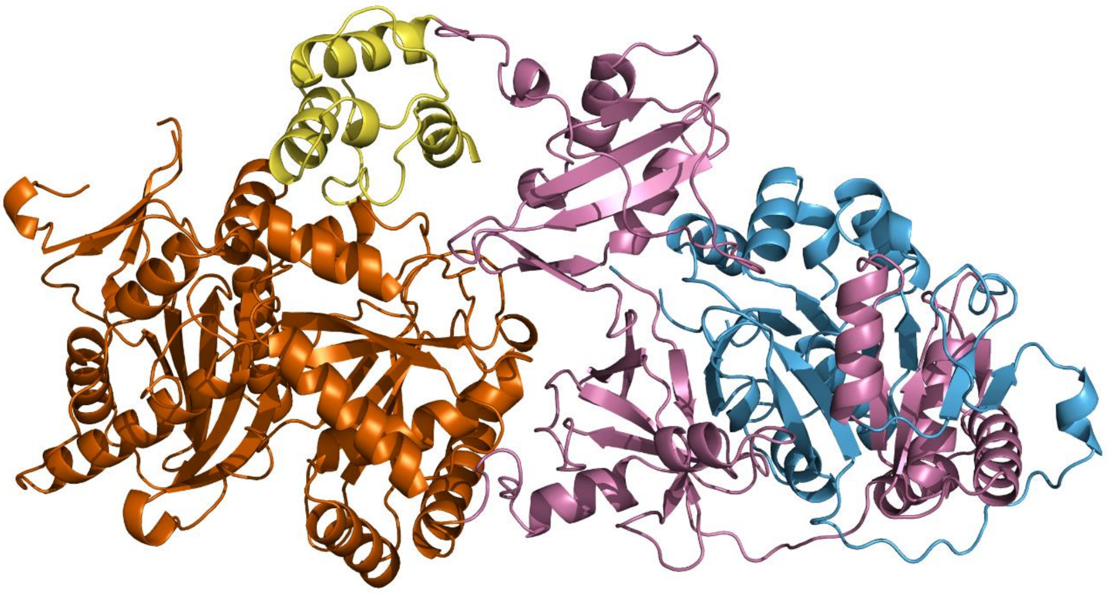

Homology model of WolG1 made with Phyre2[10] based on the structure of surfactin synthetase (PDB: 2VSQ)[42]. The condensation and thiolation domains are coloured orange and yellow, respectively. The adenylation is coloured pink, with the recombinant region coloured in blue.

**2.12 Supplementary Figure 12:** The evolutionary relationships of the desotamides, wollamides and rimosamides.

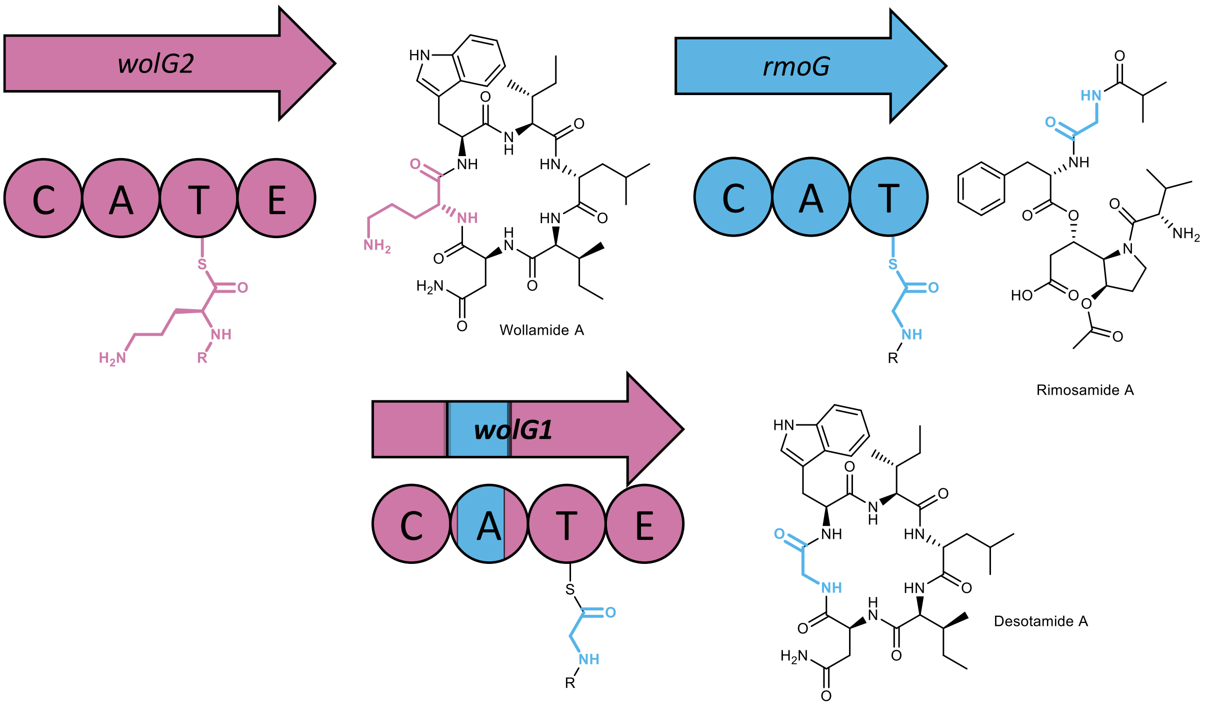

The condensation (C), adenylation (A), thiolation (T) and epimerization (E) domain architecture of the final modules of the WolG1/WolG2 NRPS assembly line enzymes are shown (R corresponds to the proceeding residues of the NRPS bound peptides).

#### 3. Supplementary Tables

**3.1 Supplementary Table 1:** Genome assembly information for Microbial Screening Technology (MST) *Streptomyces* spp. strains sequenced in this study

| | MST-7054 | MST-71321 | MST-71458 | MST-94754 | MST-110588 | MST-127221 |
|---|---|---|---|---|---|---|
| **Sequencing Platform** | Illumina | Illumina | Illumina | Illumina | PacBio | Illumina |
| **Total Contigs** | 4 | 21 | 4 | 6 | 1 | 5 |
| **Mean Contig Length** | 1,801,242 | 453,570 | 2,322,188 | 1,486,346 | 7,915,503 | 1,780,776 |
| **N50** | 9,114,431 | 1,425,667 | 9,288,751 | 6,331,616 | 7,915,503 | 8,773,046 |
| **Total Bases** | 9,114,431 | 9,524,967 | 9,288,751 | 8,918,075 | 7,915,503 | 8,903,878 |

**3.2 Supplementary Table 2:** BLAST analysis of wollamide (*wol*) BGC from *Streptomyces* sp. MST-110588

| Name | Start Position | End Position | Residues | Direction | Top Hit | Identity (%) | Coverage (%) | Role |
|---|---|---|---|---|---|---|---|---|
| Orf-1 | 4,622,717 | 4,623,274 | 345 | reverse | - | - | - | - |
| Orf-2 | 4,623,231 | 4,624,019 | 186 | forward | ABC transporter [*Streptomyces* sp. KCB13F003] | 63 | 95 | Transport |
| WolP | 4,624,440 | 4,624,853 | 263 | reverse | thioesterase [*Streptomyces* sp. 769] | 69 | 90 | Thioesterase |
| WolO | 4,625,211 | 4,625,879 | 138 | forward | hypothetical protein [*Streptomyces* sp. L-9-10] | 52 | 96 | Unknown |
| WolN | 4,625,867 | 4,627,078 | 223 | reverse | LuxR family transcriptional regulator [*Streptomyces* sp. KCB13F003] | 95 | 99 | Regulator |
| WolM | 4,627,420 | 4,628,238 | 404 | reverse | sensor histidine kinase [*Streptomyces* sp. L-9-10] | 82 | 99 | Regulator |
| WolL | 4,628,235 | 4,631,006 | 273 | forward | putative ABC transport system ATP-binding protein [*Streptomyces yunnanensis*] | 79 | 99 | Transport |
| WolK | 4,631,092 | 4,631,511 | 924 | forward | ABC transporter permease [*Streptomyces* sp. KCB13F003] | 71 | 98 | Transport |
| WolX | 4,631,762 | 4,633,141 | 140 | forward | hypothetical protein [*Streptomyces* sp. KCB13F003] | 59 | 94 | Unknown |
| WolJ | 4,633,374 | 4,637,105 | 460 | forward | DsaJ [*Streptomyces scopuliridis*] | 68 | 99 | Peptide Cyclase |
| WolI | 4,637,078 | 4,647,664 | 1244 | forward | amino acid adenylation domain-containing protein [*Streptomyces* sp. L-9-10] | 71 | 91 | NRPS |
| WolH | 4,647,843 | 4,655,768 | 3529 | forward | NRPS [*Streptomyces* sp. KCB13F003] | 67 | 99 | NRPS |
| WolG2 | 4,655,877 | 4,656,086 | 2642 | forward | DsaG [*Streptomyces scopuliridis*] | 66 | 99 | NRPS |
| WolF2 | 4,656,171 | 4,656,551 | 70 | forward | MbtH protein [*Streptomyces* sp. KCB13F003] | 87 | 98 | NRPS |
| WolE | 4,656,623 | 4,664,527 | 127 | forward | nuclear transport factor 2 family protein [*Streptomyces* sp. L-9-10] | 79 | 99 | Amino acid biosynthesis |
| WolG1 | 4,664,584 | 4,664,835 | 2635 | forward | DsaG [*Streptomyces scopuliridis*] | 72 | 99 | NRPS |
| WolF1 | 4,665,154 | 4,666,323 | 84 | forward | MbtH protein [*Streptomyces* sp. KCB13F003] | 61 | 76 | NRPS |
| WolD | 4,666,320 | 4,666,823 | 390 | forward | DsaD [*Streptomyces scopuliridis*] | 82 | 94 | Amino acid biosynthesis |
| WolC | 4,667,176 | 4,668,030 | 168 | forward | YbaK/prolyl-tRNA synthetase associated domain-containing protein [*Streptomyces* sp. L-9-10] | 84 | 98 | Unknown |
| WolW | 4,668,143 | 4,668,934 | 285 | forward | amidinotransferase [*Streptomyces* sp. KCB13F003] | 91 | 93 | Amino acid biosynthesis |
| WolB | 4,669,242 | 4,670,300 | 264 | forward | indole-3-glycerol phosphate synthase TrpC [*Streptomyces* sp. NRRL S-1813] | 76 | 99 | Amino acid biosynthesis |
| WolV | 4,622,717 | 4,623,274 | 353 | reverse | anthranilate phosphoribosyltransferase [*Streptomyces decoyicus*] | 83 | 99 | Amino acid biosynthesis |

| Name | Start Position | End Positon | Residues | Direction | Top Hit | Identity (%) | Coverage (%) | Role |
|---|---|---|---|---|---|---|---|---|
| WolU | 4,670,408 | 4,671,829 | 474 | forward | PLP-dependent aminotransferase family protein [*Streptomyces decoyicus*] | 82 | 99 | Amino acid biosynthesis |
| WolT | 4,671,909 | 4,672,484 | 192 | reverse | aminodeoxychorismate/anthranilate synthase component II [*Streptomyces decoyicus*] | 85 | 97 | Amino acid biosynthesis |
| WolS | 4,672,481 | 4,674,127 | 549 | reverse | anthranilate synthase component I family protein [*Streptomyces* sp. NRRL S-1813] | 69 | 99 | Amino acid biosynthesis |
| WolR | 4,674,491 | 4,675,834 | 448 | forward | phospho-2-dehydro-3-deoxyheptonate aldolase [*Streptomyces* sp. KCB13F003] | 85 | 97 | Amino acid biosynthesis |
| WolA | 4,676,859 | 4,677,629 | 257 | forward | DsaA [*Streptomyces scopuliridis*] | 76 | 99 | Regulator |
| Orf+1 | 4,678,250 | 4,678,567 | 106 | reverse | DUF4190 domain-containing protein [*Streptomyces varsoviensis*] | 70 | 79 | - |
| Orf+2 | 4,678,564 | 4,678,728 | 55 | reverse | hypothetical protein [*Streptomyces albospinus*] | 74 | 98 | - |
| Orf+3 | 4,678,852 | 4,680,309 | 486 | reverse | M6 family metalloprotease domain-containing protein [Streptomyces sp. ID38640] | 65 | 79 | - |

Supplementary Table 2 continued.

**3.3 Supplementary Table 3:** BLAST analysis of desotamide (*dsa*) BGC from *Streptomyces* sp. MST-70754

| Name | Start Position | End Positon | Residues | Direction | Top Hit | Identity (%) | Coverage (%) |
|---|---|---|---|---|---|---|---|
| Orf-6 | 1 | 984 | 328 | forward | hypothetical protein [*Streptomyces scopuliridis*] | 89 | 99 |
| Orf-5 | 1113 | 3332 | 740 | reverse | NADP-dependent isocitrate dehydrogenase [*Streptomyces* sp. L-9-10] | 98 | 99 |
| Orf-4 | 3435 | 3821 | 129 | reverse | D-ribose pyranase [*Streptomyces scopuliridis*] | 93 | 99 |
| Orf-3 | 3818 | 4798 | 327 | reverse | ribokinase [*Streptomyces scopuliridis*] | 89 | 99 |
| Orf-2 | 4848 | 6824 | 659 | reverse | inner-membrane translocator [*Streptomyces scopuliridis*] | 99 | 99 |
| Orf-1 | 6814 | 8370 | 519 | reverse | ATP-binding protein [*Streptomyces scopuliridis*] | 99 | 99 |
| DsaQ | 8367 | 9506 | 380 | reverse | DsaQ [*Streptomyces scopuliridis*] | 98 | 99 |
| DsaX | 9997 | 10614 | 206 | forward | hypothetical protein [*Streptomyces* sp. L-9-10] | 77 | 77 |
| DsaP | 10517 | 11188 | 224 | reverse | DsaP [*Streptomyces scopuliridis*] | 97 | 99 |
| DsaO | 11642 | 12091 | 150 | forward | DsaO [*Streptomyces scopuliridis*] | 98 | 99 |
| DsaN | 12264 | 12932 | 223 | reverse | response regulator transcription factor [*Streptomyces* sp. L-9-10] | 98 | 99 |
| DsaM | 12920 | 14131 | 404 | reverse | DsaM [*Streptomyces scopuliridis*] | 98 | 99 |
| DsaL | 14411 | 15229 | 273 | forward | ABC transporter ATP-binding protein [*Streptomyces* sp. L-9-10] | 93 | 99 |
| DsaK | 15235 | 17994 | 920 | forward | DsaK [*Streptomyces scopuliridis*] | 99 | 99 |
| DsaJ | 18164 | 19573 | 470 | forward | DsaJ [*Streptomyces scopuliridis*] | 99 | 99 |
| DsaH | 19805 | 34177 | 4791 | forward | NRPS [*Streptomyces* sp. KCB13F003] | 65 | 99 |
| DsaG | 34298 | 42241 | 2648 | forward | DsaG [*Streptomyces scopuliridis*] | 97 | 99 |
| DsaF | 42320 | 42529 | 70 | forward | DsaF [*Streptomyces scopuliridis*] | 100 | 98 |
| DsaE | 42618 | 42992 | 125 | forward | DsaE [*Streptomyces scopuliridis*] | 98 | 99 |
| DsaD | 43122 | 44255 | 378 | forward | DsaD [*Streptomyces scopuliridis*] | 98 | 99 |
| DsaC | 44252 | 44755 | 168 | forward | DsaC [*Streptomyces scopuliridis*] | 99 | 99 |
| DsaB | 44826 | 45638 | 271 | forward | DsaB [*Streptomyces scopuliridis*] | 98 | 99 |
| DsaY | 46439 | 46624 | 62 | forward | - | - | - |
| DsaA | 46599 | 47516 | 306 | forward | DsaA [*Streptomyces scopuliridis*] | 99 | 99 |
| Orf+1 | 47650 | 48351 | 234 | reverse | hydrolase [*Streptomyces scopuliridis*] | 98 | 99 |
| Orf+2 | 48365 | 48847 | 161 | reverse | transcriptional regulator [*Streptomyces scopuliridis*] | 100 | 99 |
| Orf+3 | 48905 | 50305 | 467 | reverse | hypothetical protein [*Streptomyces scopuliridis*] | 89 | 99 |

**3.4 Supplementary Table 4:** BLAST analysis of desotamide (*dsa*) BGC from *Streptomyces* sp. MST-71321

| Name | Start Position | End Positon | Residues | Direction | Top Hit | Identity (%) | Coverage (%) |
|---|---|---|---|---|---|---|---|
| Orf-1 | 1 | 618 | 206 | forward | hypothetical protein [*Streptomyces* sp. L-9-10] | 77 | 77 |
| DsaP | 521 | 1192 | 224 | reverse | DsaP [*Streptomyces scopuliridis*] | 97 | 99 |
| DsaO | 1655 | 2104 | 150 | forward | DsaO [*Streptomyces scopuliridis*] | 97 | 99 |
| DsaN | 2277 | 2945 | 223 | reverse | response regulator transcription factor [*Streptomyces* sp. L-9-10] | 98 | 99 |
| DsaM | 2933 | 4144 | 404 | reverse | DsaM [*Streptomyces scopuliridis*] | 99 | 99 |
| DsaL | 4424 | 5242 | 273 | forward | ABC transporter ATP-binding protein [*Streptomyces* sp. L-9-10] | 93 | 99 |
| DsaK | 5248 | 8007 | 920 | forward | DsaK [*Streptomyces scopuliridis*] | 99 | 99 |
| DsaJ | 8177 | 9586 | 470 | forward | DsaJ [*Streptomyces scopuliridis*] | 99 | 99 |
| DsaH | 9818 | 24172 | 4785 | forward | NRPS [*Streptomyces* sp. KCB13F003] | 65 | 99 |
| DsaG | 24296 | 32239 | 2648 | forward | DsaG [*Streptomyces scopuliridis*] | 97 | 99 |
| DsaF | 32324 | 32533 | 70 | forward | DsaF [*Streptomyces scopuliridis*] | 100 | 98 |
| DsaE | 32603 | 32977 | 125 | forward | DsaE [*Streptomyces scopuliridis*] | 98 | 99 |
| DsaD | 33110 | 34243 | 378 | forward | DsaD [*Streptomyces scopuliridis*] | 98 | 99 |
| DsaC | 34240 | 34743 | 168 | forward | DsaC [*Streptomyces scopuliridis*] | 99 | 99 |
| DsaB | 34813 | 35625 | 271 | forward | DsaB [*Streptomyces scopuliridis*] | 97 | 99 |
| DsaY | 36438 | 36623 | 62 | forward | - | - | - |
| DsaA | 36598 | 37515 | 306 | forward | DsaA [*Streptomyces scopuliridis*] | 99 | 99 |
| Orf+1 | 37641 | 38342 | 234 | reverse | hydrolase [*Streptomyces scopuliridis*] | 98 | 99 |
| Orf+2 | 38356 | 38838 | 161 | reverse | transcriptional regulator [*Streptomyces scopuliridis*] | 100 | 99 |
| Orf+3 | 38896 | 40293 | 466 | reverse | hypothetical protein [*Streptomyces scopuliridis*] | 90 | 99 |

**3.5 Supplementary Table 5:** BLAST analysis of desotamide (*dsa*) BGC from *Streptomyces* sp. MST-71458

| Name | Start Position | End Positon | Residues | Direction | Top Hit | Identity (%) | Coverage (%) |
|---|---|---|---|---|---|---|---|
| Orf-1 | 1 | 639 | 213 | forward | hypothetical protein [*Streptomyces* sp. L-9-10] | 77 | 78 |
| DsaP | 521 | 1213 | 231 | reverse | DsaP [*Streptomyces scopuliridis*] | 92 | 100 |
| DsaO | 1667 | 2116 | 150 | forward | DsaO [*Streptomyces scopuliridis*] | 97 | 100 |
| DsaN | 2290 | 2958 | 223 | reverse | response regulator transcription factor [*Streptomyces* sp. L-9-10] | 98 | 100 |
| DsaM | 2946 | 4157 | 404 | reverse | DsaM [*Streptomyces scopuliridis*] | 98 | 100 |
| DsaL | 4437 | 5255 | 273 | forward | ABC transporter ATP-binding protein [*Streptomyces* sp. L-9-10] | 92 | 99 |
| DsaK | 5261 | 8020 | 920 | forward | DsaK [*Streptomyces scopuliridis*] | 98 | 100 |
| DsaJ | 8190 | 9599 | 470 | forward | DsaJ [*Streptomyces scopuliridis*] | 99 | 100 |
| DsaH | 9830 | 24238 | 4803 | forward | NRPS [*Streptomyces* sp. KCB13F003] | 65 | 100 |
| DsaG | 24343 | 32280 | 2646 | forward | DsaG [*Streptomyces scopuliridis*] | 97 | 100 |
| DsaF | 32347 | 32556 | 70 | forward | DsaF [*Streptomyces scopuliridis*] | 100 | 100 |
| DsaE | 32662 | 33036 | 125 | forward | DsaE [*Streptomyces scopuliridis*] | 99 | 100 |
| DsaD | 33125 | 34258 | 378 | forward | DsaD [*Streptomyces scopuliridis*] | 97 | 100 |
| DsaC | 34255 | 34758 | 168 | forward | DsaC [*Streptomyces scopuliridis*] | 99 | 100 |
| DsaB | 34828 | 35640 | 271 | forward | DsaB [*Streptomyces scopuliridis*] | 99 | 100 |
| DsaA | 36620 | 37537 | 306 | forward | DsaA [*Streptomyces scopuliridis*] | 100 | 100 |
| Orf+1 | 37646 | 38347 | 234 | reverse | hydrolase [*Streptomyces scopuliridis*] | 98 | 100 |
| Orf+2 | 38361 | 38843 | 161 | reverse | winged helix-turn-helix domain-containing protein [*Streptomyces scopuliridis*] | 90 | 100 |
| Orf+3 | 38901 | 40286 | 462 | reverse | FUSC family protein [*Streptomyces scopuliridis*] | 92 | 98 |

**3.6 Supplementary Table 6:** BLAST analysis of desotamide (*dsa*) BGC from *Streptomyces* sp. MST-94754

| Name | Start Position | End Positon | Residues | Direction | Top Hit | Identity (%) | Coverage (%) |
|---|---|---|---|---|---|---|---|
| Orf-1 | 1 | 639 | 213 | forward | hypothetical protein [*Streptomyces* sp. L-9-10] | 92 | 77 |
| DsaP | 521 | 1219 | 233 | reverse | thioesterase [*Streptomyces* sp. L-9-10] | 91 | 99 |
| DsaO | 1673 | 2122 | 150 | forward | hypothetical protein [*Streptomyces* sp. L-9-10] | 95 | 99 |
| DsaN | 2340 | 3008 | 223 | reverse | response regulator transcription factor [*Streptomyces* sp. L-9-10] | 100 | 99 |
| DsaM | 2996 | 4207 | 404 | reverse | DsaM [*Streptomyces scopuliridis*] | 98 | 99 |
| DsaL | 4494 | 5312 | 273 | forward | ABC transporter ATP-binding protein [*Streptomyces* sp. L-9-10] | 99 | 99 |
| DsaK | 5318 | 8086 | 923 | forward | ABC transporter permease [*Streptomyces* sp. L-9-10] | 96 | 99 |
| DsaJ | 8405 | 9814 | 470 | forward | serine hydrolase [*Streptomyces* sp. L-9-10] | 96 | 99 |
| DsaI | 10049 | 14797 | 1583 | forward | amino acid adenylation domain-containing protein [*Streptomyces* sp. L-9-10] | 90 | 84 |
| DsaH | 14854 | 24138 | 3095 | forward | DsaH [*Streptomyces scopuliridis*] | 89 | 99 |
| DsaG | 24246 | 32198 | 2651 | forward | DsaG [*Streptomyces scopuliridis*] | 89 | 99 |
| DsaF | 32238 | 32447 | 70 | forward | MbtH family protein [*Streptomyces* sp. L-9-10] | 96 | 98 |
| DsaE | 32544 | 32918 | 125 | forward | nuclear transport factor 2 family protein [*Streptomyces* sp. L-9-10] | 97 | 99 |
| DsaD | 32976 | 34133 | 386 | forward | branched-chain amino acid aminotransferase [*Streptomyces* sp. L-9-10] | 93 | 99 |
| DsaC | 34130 | 34633 | 168 | forward | DsaC [*Streptomyces scopuliridis*] | 94 | 99 |
| DsaB | 34705 | 35520 | 272 | forward | indole-3-glycerol phosphate synthase TrpC [*Streptomyces* sp. L-9-10] | 93 | 92 |
| DsaR | 35577 | 36182 | 202 | reverse | TetR family transcriptional regulator [*Streptomyces* sp. NEAU-C40] | 85 | 94 |
| DsaS | 36314 | 37213 | 300 | forward | SDR family oxidoreductase [*Streptomyces* sp. NEAU-C40] | 88 | 99 |
| DsaA | 38559 | 39476 | 306 | forward | DsaA [*Streptomyces scopuliridis*] | 93 | 99 |
| Orf+1 | 39669 | 40370 | 234 | reverse | HAD family phosphatase [*Streptomyces* sp. L-9-10] | 97 | 99 |
| Orf+2 | 40384 | 40866 | 161 | reverse | Lrp/AsnC family transcriptional regulator [*Streptomyces* sp. L-9-10] | 99 | 99 |
| Orf+3 | 40938 | 42365 | 476 | reverse | hypothetical protein [*Streptomyces* sp. L-9-10] | 84 | 99 |

**3.7 Supplementary Table 7:** BLAST analysis of desotamide (*dsa*) BGC from *Streptomyces* sp. MST-127221

| Name | Start Position | End Positon | Residues | Direction | Top Hit | Identity (%) | Coverage (%) |
|---|---|---|---|---|---|---|---|
| Orf-1 | 1 | 633 | 211 | forward | hypothetical protein [*Streptomyces* sp. L-9-10] | 77 | 93 |
| DsaP | 521 | 1213 | 231 | reverse | thioesterase [*Streptomyces* sp. L-9-10] | 94 | 99 |
| DsaO | 1666 | 2115 | 150 | forward | hypothetical protein [*Streptomyces* sp. L-9-10] | 96 | 99 |
| DsaN | 2305 | 2973 | 223 | reverse | response regulator transcription factor [*Streptomyces* sp. L-9-10] | 100 | 99 |
| DsaM | 2961 | 4172 | 404 | reverse | sensor histidine kinase [*Streptomyces* sp. L-9-10] | 98 | 99 |
| DsaL | 4459 | 5277 | 273 | forward | ABC transporter ATP-binding protein [*Streptomyces* sp. L-9-10] | 100 | 99 |
| DsaK | 5283 | 8051 | 923 | forward | ABC transporter permease [*Streptomyces* sp. L-9-10] | 98 | 99 |
| DsaJ | 8370 | 9779 | 470 | forward | serine hydrolase [*Streptomyces* sp. L-9-10] | 97 | 99 |
| DsaH | 9924 | 24395 | 4824 | forward | NRPS [*Streptomyces* sp. KCB13F003] | 66 | 99 |
| DsaG | 24509 | 32494 | 2662 | forward | DsaG [*Streptomyces scopuliridis*] | 89 | 99 |
| DsaF | 32532 | 32741 | 70 | forward | MbtH family protein [*Streptomyces* sp. L-9-10] | 100 | 98 |
| DsaE | 32802 | 33176 | 125 | forward | nuclear transport factor 2 family protein [*Streptomyces* sp. L-9-10] | 100 | 99 |
| DsaD | 33234 | 34400 | 389 | forward | branched-chain amino acid aminotransferase [*Streptomyces* sp. L-9-10] | 94 | 99 |
| DsaC | 34397 | 34900 | 168 | forward | YbaK/prolyl-tRNA synthetase associated domain-containing protein [*Streptomyces* sp. L-9-10] | 97 | 99 |
| DsaB | 34987 | 35793 | 269 | forward | indole-3-glycerol phosphate synthase TrpC [*Streptomyces* sp. L-9-10] | 97 | 99 |
| DsaT | 35821 | 36393 | 191 | reverse | TetR family transcriptional regulator [*Streptomyces* sp. NEAU-C40] | 82 | 99 |
| DsaS | 36516 | 37412 | 299 | forward | SDR family oxidoreductase [*Streptomyces* sp. NEAU-C40] | 88 | 99 |
| DsaY | 37630 | 38028 | 133 | reverse | - | - | - |
| DsaA | 38751 | 39668 | 306 | forward | DsaA [*Streptomyces scopuliridis*] | 94 | 99 |
| Orf+1 | 39709 | 40410 | 234 | reverse | HAD family phosphatase [*Streptomyces* sp. L-9-10] | 97 | 99 |
| Orf+2 | 40424 | 40906 | 161 | reverse | Lrp/AsnC family transcriptional regulator [*Streptomyces* sp. L-9-10] | 99 | 99 |
| Orf+3 | 40964 | 42394 | 477 | reverse | hypothetical protein [*Streptomyces* sp. L-9-10] | 90 | 99 |

**3.8 Supplementary Table 8:** Adenylation Domain Specificity Codes of the Wollamide (*wol*) BGC and RmoG (Orf6595)

| A Domain | 235 | 236 | 239 | 278 | 299 | 301 | 322 | 330 | 331 | 517 | Observed Substrate |
|---|---|---|---|---|---|---|---|---|---|---|---|
| 1AMU | D | A | W | T | I | A | A | I | C | K | L-phenylalanine |
| WolI A | D | V | A | M | T | G | M | V | T | K | L-tryptophan |
| WolH A1 | D | A | L | F | V | A | A | V | A | K | L-isoleucine |
| WolH A2 | D | A | L | F | V | A | A | V | V | K | L-leucine |
| WolH A3 | D | A | L | W | S | G | G | V | F | K | L-leucine |
| WolG2 A1 | D | L | T | K | G | E | E | V | G | K | L-asparagine |
| WolG2 A2 | S | F | S | D | L | G | F | V | D | K | L-ornithine |
| WolG1 A1 | D | L | T | K | G | E | E | V | G | K | L-asparagine |
| WolG1 A2 | D | I | L | Q | G | E | L | I | W | K | glycine |
| RmoG A | D | I | L | Q | L | G | V | I | W | K | glycine |

Substrate specificity codes of each adenylation domain were identified by alignment with PheA from *Bacillus brevis* (1AMU).

**3.9 Supplementary Table 9:** RDP analysis of all NRPS encoded adenylation domains in the *Streptomyces* sp. MST-110588 genome

| Recombinant | Major parent | Minor parent | Recombinant region | R | G | B | M | C | S | T |
|---|---|---|---|---|---|---|---|---|---|---|
| ***<u>wolG1A2</u>*** | ***wolG2A2*** | ***orf6595A*** | **Y116-G343** | **+** | **+** | **+** | **+** | **+** | **+** | **+** |
| *orf1237A* | Unknown | *orf1236A* | I352-I448 | + | + | + | + | + | + | + |
| *orf6593A2* | *orf1236A* | Unknown | M139-N354 | + | - | + | - | + | + | - |
| *orf1237A* | *orf6652A* | Unknown | K223-M241 | + | + | - | + | - | - | - |
| *orf3282A* | *orf6595A* | Unknown | I357-H452 | - | - | + | + | - | + | - |
| *wolHA1* | *wolHA3* | *orf1227A* | N219-K316 | - | - | - | + | + | + | - |
| *wolHA2* | *wolHA3* | *orf1227A* | N219-K316 | - | - | - | + | + | + | - |
| *wolG1A1* | *wolHA3* | *orf1227A* | N219-K316 | - | - | - | + | + | + | - |
| *wolG2A1* | *wolHA3* | *orf1227A* | N219-K316 | - | - | - | + | + | + | - |
| *orf6593A1* | *orf6652A* | Unknown | K314-R389 | + | - | - | - | - | + | - |
| *orf252A* | *orf253A* | *orf6595A* | T173-L202 | - | - | - | + | + | - | - |
| *orf363A* | *wolG2A2* | *orf6625* | L91-L450 | - | - | - | + | + | - | - |
| *orf1224A* | *orf6593A1* | *orf289A* | K84-A332 | - | - | - | + | + | - | - |
| *orf1236A* | Unknown | *orf6594* | M95-I377 | - | - | - | + | - | + | - |

Predicted recombinants from RDP4 for which breakpoint analysis could be applied. Recombination region refers to the approximate homologous positions in the L-phenylalanine activating domain of gramicidin synthase (1AMU). Events are sorted by the whether or not (+/-) the event was predicted by a given detection method (R: RDP[13]; G: GENECONV[14]; B: BootScan[15]; M: MaxChi[16]; C: Chimaera[17]; S: SiScan[18]; L: LARD[19]).

**3.10 Supplementary Table S10:** cblaster analysis of minor parent BGC from *Streptomyces* sp. MST-110588

| Organism | MIBIG Accession | Product | 6603 | 02 | 01 | 00 | 99 | E | F | 96 | G | H | I | 92 | J | K | L | M | 87 | 86 | 6585 |
|---|---|---|---|---|---|---|---|---|---|---|---|---|---|---|---|---|---|---|---|---|---|
| *Streptomyces rimosus* subsp. rimosus ATCC 10970 | BGC0001760 | Rimosamide | 0 | 0 | 0 | 0 | 0 | 1 | 1 | 0 | 1 | 1 | 1 | 1 | 1 | 1 | 1 | 1 | 0 | 0 | 0 |
| *Streptomyces canus* ATCC 12647 | BGC0001406 | Telomycin | 1 | 1 | 0 | 0 | 0 | 0 | 0 | 0 | 1 | 0 | 1 | 0 | 0 | 0 | 0 | 0 | 0 | 0 | 0 |
| *Streptomyces* sp. SANK 62799 SANK 62799 | BGC0000288 | A-503083 | 0 | 0 | 0 | 0 | 0 | 0 | 0 | 0 | 0 | 0 | 0 | 0 | 0 | 0 | 0 | 0 | 1 | 1 | 1 |

Results from a local cblaster[43] search querying the genes surrounding *orf6595* against a local copy of the MIBiG[44] database (16/10/19). Genes are named according to their *rmo* homologue or position in the *S.* sp MST115088 genome where no homologue is avaliable. The minor parent involved in the recombination event discussed in this paper, *rmoG* (*orf6595*), is underlined. Cells are shaded to reflect the number of hits against each cluster.

**3.11 Supplementary Table S11:** Hydroxylamine trapping assay of adenylation domain specificity

| Amino Acid | WolG2A2 | | WolG1A2 | | RmoG | | Recombinant | |
|---|---|---|---|---|---|---|---|---|
| | A545 | SE | A545 | SE | A545 | SE | A545 | SE |
| A | 0.00 | 3.E-04 | 0.00 | 8.E-04 | 0.00 | 6.E-04 | 0.01 | 7.E-04 |
| C | 0.00 | 3.E-03 | 0.00 | 1.E-03 | 0.00 | 3.E-04 | 0.01 | 6.E-04 |
| D | 0.01 | 1.E-03 | 0.00 | 2.E-03 | 0.00 | 6.E-04 | 0.01 | 7.E-04 |
| E | 0.00 | 3.E-04 | 0.00 | 6.E-04 | 0.00 | 6.E-04 | 0.01 | 1.E-03 |
| F | 0.00 | 7.E-04 | 0.00 | 4.E-04 | 0.00 | 2.E-03 | 0.01 | 7.E-04 |
| G | 0.00 | 6.E-04 | 0.02 | 7.E-03 | 0.03 | 7.E-03 | 0.07 | 3.E-03 |
| H | 0.00 | 1.E-03 | 0.00 | 1.E-03 | 0.00 | 1.E-03 | 0.01 | 3.E-04 |
| I | 0.00 | 7.E-04 | 0.00 | 5.E-04 | 0.00 | 9.E-04 | 0.01 | 3.E-04 |
| K | 0.00 | 3.E-04 | 0.00 | 1.E-04 | 0.00 | 1.E-03 | 0.01 | 3.E-04 |
| L | 0.00 | 6.E-04 | 0.00 | 6.E-04 | 0.00 | 3.E-04 | 0.01 | 9.E-04 |
| M | 0.00 | 3.E-04 | 0.00 | 3.E-04 | 0.00 | 3.E-04 | 0.01 | 3.E-04 |
| N | 0.01 | 6.E-04 | 0.00 | 2.E-03 | 0.00 | 3.E-03 | 0.01 | 6.E-04 |
| O | 0.02 | 2.E-03 | 0.00 | 4.E-03 | 0.00 | 7.E-04 | 0.01 | 9.E-04 |
| P | 0.00 | 3.E-04 | 0.00 | 1.E-04 | 0.00 | 1.E-03 | 0.01 | 0.E00 |
| Q | 0.00 | 1.E-03 | 0.00 | 1.E-03 | 0.00 | 2.E-03 | 0.01 | 3.E-04 |
| R | 0.00 | 1.E-03 | 0.00 | 5.E-04 | 0.00 | 1.E-03 | 0.01 | 6.E-04 |
| S | 0.00 | 9.E-04 | 0.00 | 1.E-03 | 0.00 | 3.E-04 | 0.01 | 9.E-04 |
| T | 0.00 | 3.E-04 | 0.00 | 1.E-04 | 0.00 | 6.E-04 | 0.01 | 7.E-04 |
| V | 0.00 | 2.E-03 | 0.00 | 7.E-04 | 0.00 | 9.E-04 | 0.01 | 2.E-03 |
| W | 0.00 | 9.E-04 | 0.00 | 2.E-04 | 0.00 | 7.E-04 | 0.01 | 1.E-03 |
| Y | 0.00 | 9.E-04 | 0.00 | 6.E-04 | 0.00 | 9.E-04 | 0.01 | 6.E-04 |

Each domain was tested against the twenty proteinogenic amino acids and L-ornithine (O). Average absorbance at 545 nm measured during the assay across at least three replicates per sample and standard errors are displayed. Absorption values are coloured according to intensity.

**3.12 Supplementary Table 12:** Strains used in this study

| Strain | Description | Reference |
|---|---|---|
| *Escherichia coli* ET12567/pUZ8002 | Demethylating conjugative strain containing the helper plasmid pUZ8002. | [27] |
| *Escherichia coli* ET12567/pUZ8002 pBO1 | Conjugation of the plasmid pBO1 to *Streptomyces* strains. | This study |
| *Escherichia coli* ET12567/pUZ8002 pBO1-*wolG2* | Conjugation of the plasmid pBO1-*wolG2* to *Streptomyces*. | This study |
| *Escherichia coli* NiCo21 (DE3) | Protein expression strain. | |
| *Escherichia coli* NiCo21 (DE3)/pET28a(+)-*orf6595* | Expression of the adenylation domain ORF6595. | This study |
| *Escherichia coli* NiCo21 (DE3)/pET28a(+)-*orf6595*; pCDFDuet-1-*wolF2* | Coexpression of the adenylation domain ORF6595 and the MbtH-like protein WolF2. | This study |
| *Escherichia coli* NiCo21 (DE3)/pET28a(+)-*wolG1A2* | Expression of the adenylation domain WolG1A2. | This study |
| *Escherichia coli* NiCo21 (DE3)/pET28a(+)-*wolG1A2*; pCDFDuet-1-*wolF2* | Coexpression of the adenylation domain WolG1A2 and the MbtH-like protein WolF2. | This study |
| *Escherichia coli* NiCo21 (DE3)/pET28a(+)-*wolG2A2* | Expression of the adenylation domain WolG2A2. | This study |
| *Escherichia coli* NiCo21 (DE3)/pET28a(+)-*wolG2A2*; pCDFDuet-1-*wolF2* | Coexpression of the adenylation domain WolG2A2 and the MbtH-like protein WolF2. | This study |
| *Escherichia coli* NiCo21 (DE3)/pET28a(+)-*wolG2MP* | Expression of the adenylation domain WolG2A2::RmoG. | This study |
| *Escherichia coli* NiCo21 (DE3)/pET28a(+)-*wolG2MP*; pCDFDuet-1-*wolF2* | Coexpression of the hybrid adenylation domain WolG2A2::RmoG and the MbtH-like protein WolF2. | This study |
| *Escherichia coli* DH5α | Cloning and propogation of vectors. | [45] |
| *Escherichia coli* DH10B | F_ mcrA (mrr-hsdRMS-mcrBC), 80lacZΔ, M15, ΔlacX74 recA1 endA1 araD 139 Δ(ara, leu)7697 galU galK λrpsL (Strr) nupG | [45] |
| *Saccharomyces cerevisiae* CEN.PK 2-1C | Assembly and propogation of pBO1 and its derrivatives. | [46] |
| *Streptomyces* sp. MST-110588 | Desotamide and wollamide producing strain. | This study |
| *Streptomyces* sp. MST-127221 | Desotamide producing strain. | This study |
| *Streptomyces* sp. MST-70754 | Desotamide producing strain. | This study |
| *Streptomyces* sp. MST-7075/pBO1 | Desotamide producing strain containing empty pBO1 (control). | This study |
| *Streptomyces* sp. MST-7075/pBO1-*wolG2* | Desotamide producing strain engineered to produce wollamides through over expression of *wolG2* | This study |
| *Streptomyces* sp. MST-71321 | Desotamide producing strain. | This study |
| *Streptomyces* sp. MST-71458 | Desotamide producing strain. | This study |
| *Streptomyces* sp. MST-94754 | Desotamide producing strain. | This study |

**3.13 Supplementary Table 13:** Plasmids used in this study

| Plasmid | Description | Reference |
|---|---|---|
| pGP9 | Integrative (φBT1) *Streptomyces* expression vector; propagates in E. coli for cloning/conjugation; $Apr^R$ | [23] |
| pFF62A | Yeast shuttle vector used in the construction of pBO1 | [24] |
| pUZ8002 | Non-transmissible oriT mobilizing plasmid; $Cm^R$ | [27] |
| pBO1 | Integrative (φBT1) *Streptomyces* expression vector; propagates in *E. coli* and *S. cerevisiae* for cloning and conjugation; $Apr^R$ | This study. |
| pBO1-*wolG2* | Integrative (φBT1) *Streptomyces* expression vector with *wolG2* cloned between the NdeI sites; propagates in *E. coli* and *S. cerevisiae* for cloning and conjugation; $Apr^R$ | This study. |
| pET28a | *E. coli* expression vector; $Kan^R$ | EMD Biosciences |
| pET28a-wolG1A1 | *E. coli* expression vector with adenylation domain coding region (WolG2A1) cloned between NdeI and XhoI sites; $Kan^R$ | This study. |
| pET28a-wolG2A2 | *E. coli* expression vector with adenylation domain coding region (WolG2A2) cloned between NdeI and XhoI sites; $Kan^R$ | This study. |
| pET28a-wolG2MP | *E. coli* expression vector with hybrid adenylation domain coding region (WolG2A2::RmoG). This vector is a derivative of pET28a-wolG2A2. | This study |
| pCDFDuet | *E. coli* expression vector; $Sm^R$ | Novagen (EMD Millipore) |
| pcDFDuet-*wolF2* | E. coli expression vector with *wolF* cloned between NdeI and KpnI sites; $Sm^R$ | This study. |

**3.14 Supplementary Table 14:** Primers used in this study

| Primer | Sequence | Annealing Temperature (°C) | Description |
|---|---|---|---|
| pGP9_seq_F1 | gagcggcggtcgaagggagatg | 50 | Sequencing and colony PCR of pBO1 inserts |
| pGP9_seq_R1 | cgagcgttctgaacaaatccag | | |
| pBO1_pGP9_F1 | aatttattcatatcaggattatcaataccatatttttgaaaaagccgtttctgtaa<br>tgaaggagaaaactcaccgaggcacttcctcgctcactgactcg | 70 | Amplification of pGP9 for the assembly of pBO1 |
| pBO1_pGP9_R1 | agcagcaccatatgatcacgtttttcattcggatctttaaacagtgcgctctga<br>tagctcagataatgattatccggcagcagaaccggaccatcaccaacgcgt<br>tggccgattcattaa | | |
| pBO1_pFF_F1 | agaacgctcggttgccgccgggcgttttttattggtgagaatccaagctaga<br>aatctgcattaatgaatcggccaacgcgttggtgatggtccggttctg | 70 | Amplification of pFF for the assembly of pBO1 |
| pBO1_pFF_R1 | ccgtattaccgcctttgagtgagctgataccgctcgccgcagccgaacgac<br>cgagcgcagcgagtcagtgagcgaggaagtgcctcggtgagttttctcc | | |
| pBO1_wolG2_F1 | ttcgagcctccttcgagccacggggccgacgatgacgacgaccaccgga<br>cgaacgcatcgattaattaaggaggacacatatggacgcggcggcatcac | 70 | Cloning of *wolG2* into pBO1 |
| pBO1_wolG2_R1 | | | |
| pCDF_wolF2_F1 | aagtataagaaggagatatacatatgtcgaacccgttcg | 60 | Coloning of *wolF2* into pCDFDuet-1 |
| pCDF_wolF2_R1 | ctcgagtctggtaaagaaacggtaccgtgtgtacgagtggtg | | |
| pET_wolG1_F1 | catcaccacagccaggatccgaatccgacgtacgcgcagctcaacga | 62 | Cloning of wolG1A2 into pET28a(+) |
| pET_wolG1_R1 | cttaagcattatgcggccgcaagcttttagatcacgcacgcctgggcca | | |
| pET_wolG2_F1 | catcaccacagccaggatccgaatccgaccttcgcggagctggaggc | 62 | Cloning of wolG2A2 into pET28a(+) |
| pET_wolG2_R1 | cttaagcattatgcggccgcaagcttttacaccacggcctgcgccacgt | | |
| pET_6595_F1 | catcaccacagccaggatccgaattcgacctacgccgaactggaagc | 60 | Cloning of orf6595A into pET28a(+) |
| pET_6595_R1 | cttaagcattatgcggccgcaagcttttaggcggccagctccacaccct | | |
| pET-G2MP-1 | ggccaccgacaccaccgggc | 20 | Cloning of wolG2A2::RmoG into pET-wolG2 |
| pET-G2MP-2 | ccagccccagcagttcggcg | 20 | Cloning of wolG2A2::RmoG into pET-wolG2 |
| pET-G2MP-3 | gcccggtggtgtcggtggcc | 20 | Cloning of wolG2A2::RmoG into pET-wolG2 |
| pET-G2MP-4 | cgccgaactgctggggctgg | 20 | Cloning of wolG2A2::RmoG into pET-wolG2 |

#### 4. References

(1) Kieser, T.; Bibb, M. J.; Buttner, M. J.; Chater, K. F.; Hopwood, D. A.; John Innes Foundation. *Practical Streptomyces Genetics*; John Innes Foundation, 2000.

(2) Hyatt, D.; Chen, G.-L.; LoCascio, P. F.; Land, M. L.; Larimer, F. W.; Hauser, L. J. Prodigal: Prokaryotic Gene Recognition and Translation Initiation Site Identification. *BMC Bioinformatics* **2010**, *11* (1), 119. https://doi.org/10.1186/1471-2105-11-119.

(3) Blin, K.; Shaw, S.; Steinke, K.; Villebro, R.; Ziemert, N.; Lee, S. Y.; Medema, M. H.; Weber, T. AntiSMASH 5.0: Updates to the Secondary Metabolite Genome Mining Pipeline. *Nucleic Acids Res.* **2019**, *47* (W1), W81–W87.

(4) Alanjary, M.; Steinke, K.; Ziemert, N. AutoMLST: An Automated Web Server for Generating Multi-Locus Species Trees Highlighting Natural Product Potential. *Nucleic Acids Res.* **2019**, *47* (W1), W276–W282. https://doi.org/10.1093/nar/gkz282.

(5) Prieto, C.; García-estrada, C.; Lorenzana, D.; Martín, J. F. NRPSSP: Non-Ribosomal Peptide Synthase Substrate Predictor. *Bioinformatics* **2012**, *28* (3), 426–427. https://doi.org/10.1093/bioinformatics/btr659.

(6) Röttig, M.; Medema, M. H.; Blin, K.; Weber, T.; Rausch, C.; Kohlbacher, O. NRPSpredictor2--a Web Server for Predicting NRPS Adenylation Domain Specificity. *Nucleic Acids Res.* **2011**, *39* (Web Server issue), W362-7. https://doi.org/10.1093/nar/gkr323.

(7) Stachelhaus, T.; Mootz, H. D.; Marahiel, M. A. The Specificity-Conferring Code of Adenylation Domains in Nonribosomal Peptide Synthetases. *Chem. Biol.* **1999**, *6* (8), 493–505. https://doi.org/10.1016/S1074-5521(99)80082-9.

(8) Minowa, Y.; Araki, M.; Kanehisa, M. Comprehensive Analysis of Distinctive Polyketide and Nonribosomal Peptide Structural Motifs Encoded in Microbial Genomes. *J. Mol. Biol.* **2007**, *368* (5), 1500–1517. https://doi.org/10.1016/j.jmb.2007.02.099.

(9) Altschul, S. F.; Gish, W.; Miller, W.; Myers, E. W.; Lipman, D. J. Basic Local Alignment Search Tool. *J. Mol. Biol.* **1990**, *215* (3), 403–410. https://doi.org/10.1016/S0022-2836(05)80360-2.

(10) Kelley, L. A.; Mezulis, S.; Yates, C. M.; Wass, M. N.; Sternberg, M. J. E. The Phyre2 Web Portal for Protein Modeling, Prediction and Analysis. *Nat. Protoc.* **2015**, *10* (6), 845–858. https://doi.org/10.1038/nprot.2015.053.

(11) Schrödinger, L. PyMOL The PyMOL Molecular Graphics System, Version 2.0.

(12) Larkin, M. A.; Blackshields, G.; Brown, N. P.; Chenna, R.; McGettigan, P. A.; McWilliam, H.; Valentin, F.; Wallace, I. M.; Wilm, A.; Lopez, R.; et al. Clustal W and Clustal X Version 2.0. *Bioinformatics* **2007**, *23* (21), 2947–2948. https://doi.org/10.1093/bioinformatics/btm404.

(13) Martin, D.; Rybicki, E. RDP: Detection of Recombination amongst Aligned Sequences. *Bioinformatics* **2000**, *16* (6), 562–563.

(14) Padidam, M.; Sawyer, S.; Fauquet, C. M. Possible Emergence of New Geminiviruses by Frequent Recombination. *Virology* **1999**, *265* (2), 218–225. https://doi.org/10.1006/VIRO.1999.0056.

(15) Salminen, M. O.; Carr, J. K.; Burke, D. S.; Mccutchan, F. E. Identification of Breakpoints in Intergenotypic Recombinants of HIV Type 1 by Bootscanning. *AIDS Res. Hum. Retroviruses* **1995**, *11* (11), 1423–1425. https://doi.org/10.1089/aid.1995.11.1423.

(16) Smith, J. M. Analyzing the Mosaic Structure of Genes. *J. Mol. Evol.* **1992**, *34* (2), 126–129.

(17) Posada, D.; Crandall, K. A. Evaluation of Methods for Detecting Recombination from DNA Sequences: Computer Simulations. *Proc. Natl. Acad. Sci. U. S. A.* **2001**, *98* (24), 13757–13762. https://doi.org/10.1073/pnas.241370698.

(18) Gibbs, M. J.; Armstrong, J. S.; Gibbs, A. J. Sister-Scanning: A Monte Carlo Procedure for Assessing Signals in Recombinant Sequences. *Bioinformatics* **2000**, *16* (7), 573–582.

(19) Holmes, E. C.; Worobey, M.; Rambaut, A. Phylogenetic Evidence for Recombination in Dengue Virus. *Mol. Biol. Evol.* **1999**, *16* (3), 405–409. https://doi.org/10.1093/oxfordjournals.molbev.a026121.

(20) Boni, M. F.; Posada, D.; Feldman, M. W. An Exact Nonparametric Method for Inferring Mosaic Structure in Sequence Triplets. *Genetics* **2007**, *176* (2), 1035–1047. https://doi.org/10.1534/genetics.106.068874.

(21) Martin, D. P.; Murrell, B.; Golden, M.; Khoosal, A.; Muhire, B. RDP4: Detection and Analysis of Recombination Patterns in Virus Genomes. *Virus Evol.* **2015**, *1* (1), vev003. https://doi.org/10.1093/ve/vev003.

(22) Price, M. N.; Dehal, P. S.; Arkin, A. P. FastTree 2 - Approximately Maximum-Likelihood Trees for Large Alignments. *PLoS One* **2010**, *5* (3). https://doi.org/10.1371/journal.pone.0009490.

(23) Kuščer, E.; Coates, N.; Challis, I.; Gregory, M.; Wilkinson, B.; Sheridan, R.; Petkovic´*, H. P. Roles of RapH and RapG in Positive Regulation of Rapamycin Biosynthesis in Streptomyces Hygroscopicus. *J. Bacteriol.* **2007**, *189* (13), 4756–4763. https://doi.org/10.1128/JB.00129-07.

(24) Schimming, O.; Fleischhacker, F.; Nollmann, F. I.; Bode, H. B. Yeast Homologous Recombination Cloning Leading to the Novel Peptides Ambactin and Xenolindicin. *ChemBioChem* **2014**, *15* (9), 1290–1294. https://doi.org/10.1002/cbic.201402065.

(25) Gietz, R. D.; Schiestl, R. H. Quick and Easy Yeast Transformation Using the LiAc/SS Carrier DNA/PEG Method. *Nat. Protoc.* **2007**, *2* (1), 35–37. https://doi.org/10.1038/nprot.2007.14.

(26) Gietz, R. D.; Schiestl, R. H. Frozen Competent Yeast Cells That Can Be Transformed with High Efficiency Using the LiAc/SS Carrier DNA/PEG Method. *Nat. Protoc.* **2007**, *2* (1), 1–4. https://doi.org/10.1038/nprot.2007.17.

(27) Paget, M. S.; Chamberlin, L.; Atrih, A.; Foster, S. J.; Buttner, M. J. Evidence That the Extracytoplasmic Function Sigma Factor SigmaE Is Required for Normal Cell Wall Structure in Streptomyces Coelicolor A3(2). *J. Bacteriol.* **1999**, *181* (1), 204–211.

(28) Watzel, J.; Hacker, C.; Duchardt-Ferner, E.; Bode, H. B.; Wöhnert, J. A New Docking Domain Type in the Peptide-Antimicrobial-Xenorhabdus Peptide Producing Nonribosomal Peptide Synthetase from Xenorhabdus Bovienii. *ACS Chem. Biol.* **2020**, *15* (4), 982–989. https://doi.org/10.1021/acschembio.9b01022.

(29) Molecular Operating Environment (MOE). Chemical Computing Group ULC: Montreal, QC 2020.

(30) Levitt, M.; Sharon, R. Accurate Simulation of Protein Dynamics in Solution. *Proceedings of the National Academy of Sciences of the United States of America*. National Academy of Sciences 1988, pp 7557–7561. https://doi.org/10.1073/pnas.85.20.7557.

(31) Fechteler, T.; Dengler, U.; Schomburg, D. Prediction of Protein Three-Dimensional Structures in Insertion and Deletion Regions: A Procedure for Searching Data Bases of Representative Protein Fragments Using Geometric Scoring Criteria. *J. Mol. Biol.* **1995**, *253* (1), 114–131. https://doi.org/10.1006/jmbi.1995.0540.

(32) Bradford, M. M. A Rapid and Sensitive Method for the Quantitation of Microgram Quantities of Protein Utilizing the Principle of Protein-Dye Binding. *Anal. Biochem.* **1976**, *72* (1–2), 248–254. https://doi.org/10.1016/0003-2697(76)90527-3.

(33) Kadi, N.; Challis, G. L. Chapter 17 Siderophore Biosynthesis. In *Methods in enzymology*; 2009; Vol. 458, pp 431–457. https://doi.org/10.1016/S0076-6879(09)04817-4.

(34) Song, Y.; Li, Q.; Liu, X.; Chen, Y.; Zhang, Y.; Sun, A.; Zhang, W.; Zhang, J.; Ju, J. Cyclic Hexapeptides from the Deep South China Sea-Derived *Streptomyces Scopuliridis* SCSIO ZJ46 Active Against Pathogenic Gram-Positive Bacteria. *J. Nat. Prod.* **2014**, *77* (8), 1937–1941. https://doi.org/10.1021/np500399v.

(35) Kaweewan, I.; Komaki, H.; Hemmi, H.; Kodani, S. Isolation and Structure Determination of New Antibacterial Peptide Curacomycin Based on Genome Mining. *Asian J. Org. Chem.* **2017**, *6* (12), 1838–1844. https://doi.org/10.1002/ajoc.201700433.

(36) Son, S.; Hong, Y.-S.; Jang, M.; Heo, K. T.; Lee, B.; Jang, J.-P.; Kim, J.-W.; Ryoo, I.-J.; Kim, W.-G.; Ko, S.-K.; et al. Genomics-Driven Discovery of Chlorinated Cyclic Hexapeptides Ulleungmycins A and B from a Streptomyces Species. *J. Nat. Prod.* **2017**, *80* (11), 3025–3031. https://doi.org/10.1021/acs.jnatprod.7b00660.

(37) Gilchrist, C. L. M.; Chooi, Y.-H. Clinker & Clustermap.Js: Automatic Generation of Gene Cluster Comparison Figures. *bioRxiv* **2020**, 2020.11.08.370650. https://doi.org/10.1101/2020.11.08.370650.

(38) Khalil, Z. G.; Salim, A. A.; Lacey, E.; Blumenthal, A.; Capon, R. J. Wollamides: Antimycobacterial Cyclic Hexapeptides from an Australian Soil Streptomyces. *Org. Lett.* **2014**, *16* (19), 5120–5123. https://doi.org/10.1021/ol502472c.

(39) Watzel, J.; Hacker, C.; Duchardt-Ferner, E.; Bode, H. B.; Wöhnert, J. A New Docking Domain Type in the Peptide-Antimicrobial-Xenorhabdus Peptide Producing Nonribosomal Peptide Synthetase from Xenorhabdus Bovienii. *ACS Chem. Biol.* **2020**. https://doi.org/10.1021/ACSCHEMBIO.9B01022.

(40) Ziemert, N.; Podell, S.; Penn, K.; Badger, J. H.; Allen, E.; Jensen, P. R. The Natural Product Domain Seeker NaPDoS: A Phylogeny Based Bioinformatic Tool to Classify Secondary Metabolite Gene Diversity. *PLoS One* **2012**, *7* (3). https://doi.org/10.1371/journal.pone.0034064.

(41) Conti, E.; Stachelhaus, T.; Marahiel, M. A.; Brick, P. Structural Basis for the Activation of Phenylalanine in the Non-Ribosomal Biosynthesis of Gramicidin S. *EMBO J.* **1997**, *16* (14), 4174–4183.

(42) Tanovic, A.; Samel, S. A.; Essen, L.-O.; Marahiel, M. A. Crystal Structure of the Termination Module of a Nonribosomal Peptide Synthetase. *Science (80-. ).* **2008**, *321* (5889), 659–663. https://doi.org/10.1126/science.1159850.

(43) Gilchrist, C. L. M.; Booth, T. J.; Chooi, Y.-H. Cblaster: A Remote Search Tool for Rapid Identification and Visualisation of Homologous Gene Clusters. *bioRxiv* **2020**, 2020.11.08.370601. https://doi.org/10.1101/2020.11.08.370601.

(44) Medema, M. H.; Kottmann, R.; Yilmaz, P.; Cummings, M.; Biggins, J. B.; Blin, K.; De Bruijn, I.; Chooi, Y. H.; Claesen, J.; Coates, R. C.; et al. Minimum Information about a Biosynthetic Gene Cluster. *Nature Chemical Biology*. Nature Publishing Group August 18, 2015, pp 625–631. https://doi.org/10.1038/nchembio.1890.

(45) Hanahan, D. *DNA Cloning: A Practical Approach*; Glover, D. M., Ed.; IRL Press: McLean,

Virginia, 1985.

(46) Nijkamp, J. F.; van den Broek, M.; Datema, E.; de Kok, S.; Bosman, L.; Luttik, M. A.; Daran-Lapujade, P.; Vongsangnak, W.; Nielsen, J.; Heijne, W. H. M.; et al. De Novo Sequencing, Assembly and Analysis of the Genome of the Laboratory Strain Saccharomyces Cerevisiae CEN.PK113-7D, a Model for Modern Industrial Biotechnology. *Microb. Cell Fact.* **2012**, *11*. https://doi.org/10.1186/1475-2859-11-36.
